# Chromosome-scale assembly of the coral endosymbiont *Symbiodinium microadriaticum* genome provides insight into the unique biology of dinoflagellate chromosomes

**DOI:** 10.1101/2020.07.01.182477

**Authors:** Ankita Nand, Ye Zhan, Octavio R. Salazar, Manuel Aranda, Christian R. Voolstra, Job Dekker

## Abstract

Dinoflagellates are major primary producers in the world’s oceans, the cause of harmful algal blooms, and endosymbionts of marine invertebrates. Much remains to be understood about their biology including their peculiar crystalline chromosomes. Here we used Hi-C to order short read-based sub-scaffolds into 94 chromosome-scale scaffolds of the genome of the coral endosymbiont *Symbiodinium microadriaticum*. Hi-C data show that chromosomes are folded as linear rods within which loci separated by up to several Mb are highly packed. Each chromosome is composed of a series of structural domains separated by boundaries. Genes are enriched towards the ends of chromosomes and are arranged in unidirectional blocks that alternate between top and bottom strands. Strikingly, the boundaries of chromosomal domains are positioned at sites where transcription of two gene blocks converges, indicating a correlation between gene orientation, transcription and chromosome folding. Some chromosomes are enriched for genes involved in specific biological processes (e.g., photosynthesis, and nitrogen-cycling), and functionally related genes tend to co-occur at adjacent sites in the genome. All chromosomes contain several repeated segments that are enriched in mobile elements. The assembly of the *S. microadriaticum* genome and initial description of its genetic and spatial organization provide a foundation for deeper exploration of the extraordinary biology of dinoflagellates and their chromosomes.

## Introduction

Dinoflagellates are single celled marine plankton, abundant in the world’s oceans, and of great economic and ecological importance ^1^. This is due to their role as primary producers ^2^, their ability to cause harmful algal blooms ^3^, and because of the symbiotic relationships they form with a broad range of marine invertebrates ^1^. In particular, dinoflagellates in the family Symbiodiniaceae ^4^ are known for their role as intracellular symbionts of reef-building corals. Symbiodiniaceae fuel the coral’s energetic needs through the provision of photosynthates, which enables corals to build the massive three-dimensional calcium carbonate structures that provide habitat for a third of all marine species and give rise to coral reef ecosystems. In recent decades, we have witnessed unprecedented loss of coral reef cover due to local and global anthropogenic insult ^5^. Coral bleaching, i.e. the loss of Symbiodiniaceae triggered by ocean warming due to climate change, is now the main driver of coral reef degradation ^6^.

Gaining better insight into the biology of dinoflagellates is critical to conceive strategies to manage harmful algal blooms, which are projected to become more frequent and severe as a consequence of climate change ^7^, and to minimize coral reef loss ^4^. Importantly, dinoflagellates seem to defy many of the cellular features found in other eukaryotes. For instance, dinoflagellates commonly use 5-hydroxymethyluracil instead of thymidine ^8^, show a paucity of transcriptional regulation ^9–11^, exhibit broad RNA editing ^12^, and seem to have (a portion of) their genes arranged in tandem arrays ^13,14^, which may at least partially explain the pervasive gene duplication observed in in their genomes ^15^. Most interestingly, dinoflagellates fold their chromosomes in a way that is distinct from other eukaryotes and that is also distinct from prokaryotes. Dinoflagellates were until recently believed to have no histones, and their DNA was reported to be in a crystal-like state ^16,17^. More recent transcriptome studies, however, confirmed that dinoflagellates do possess histones, but lack histone H1 ^18,19^. However, only a very small fraction of the genome is nucleosomal, e.g. as shown by nuclease digestion patterns ^20^. The liquid-crystalline conformation of dinoflagellates may represent a third chromosome folding state, in addition to the typical nucleosomal form in eukaryotes and the supercoiled circular form in most bacteria. Remarkably many of the same machines that fold eukaryotic chromosomes and prokaryotic chromosomes (condensins, cohesins, topoisomerases) are all present in all three groups of organisms, yet how their chromosomes are organized appears very distinct.

For decades, dinoflagellates escaped genomic analysis due to their unusually large genomes (ranging from 1-250 Gb ^21^), prohibiting genome sequencing. With the advent of next-generation sequencing a number of Symbiodiniaceae genome sequences are now available, such as the genome of *Breviolum minutum*^22^, *Fugacium kawagutii*^23^, and *Symbiodinium microadriaticum*^15^ (among others). These genome sequences are collections of short scaffolds, but not chromosome-scale assemblies. Analyses of these draft genome sequences broadly confirmed many of the posited genetic features, besides providing further insight such as the high number of genes encoded, the pervasive presence of non-canonical splice sites, or the unidirectional alignment of genes that form cluster-like gene arrangements ^15,22^.

However, we are still missing a chromosome-scale assembly of a dinoflagellate genome, which is key to providing answers to pertinent biological questions, such as how they achieve the unusual organizational folding of their chromosomes, whether the unidirectional alignment of genes is a feature conserved across chromosomes, and whether such alignment is related to features of chromosome organization and architecture. In addition, it will be highly informative to detail the distribution of tandem gene arrays across chromosomes, and whether these clusters or genes in general are spread evenly across chromosomes or arranged in ‘hotspots’. The latter giving rise to the possibility of functionally ‘enriched’ chromosomes. To start to provide answers to these questions, we generated the first chromosome-scale assembly of the genome of the dinoflagellate *Symbiodinium microadriaticum*.

## Results

### Chromosome-scale assembly of the *Symbiodinium microadriaticum* genome

Previously we used short-read Illumina sequencing to produce a set of 9,695 scaffolds for *S. microadriaticum* ^15^. This set of scaffolds covers 808,242,489 bp (scaffold N50 = 573.5 kb, contig N50 = 34.9 kb) of the estimated 1.4 Gb genome. We employed Hi-C data, representing spatial interaction frequencies between loci genome-wide ^24^, to group, order, and orient these scaffolds to generate chromosome-scale scaffolds ^25,26^. The Hi-C-assisted assembly process is described in detail in the Methods section (Methods, Supplemental Figure S1) and is summarized in Figure 1A.

**Figure 1.**
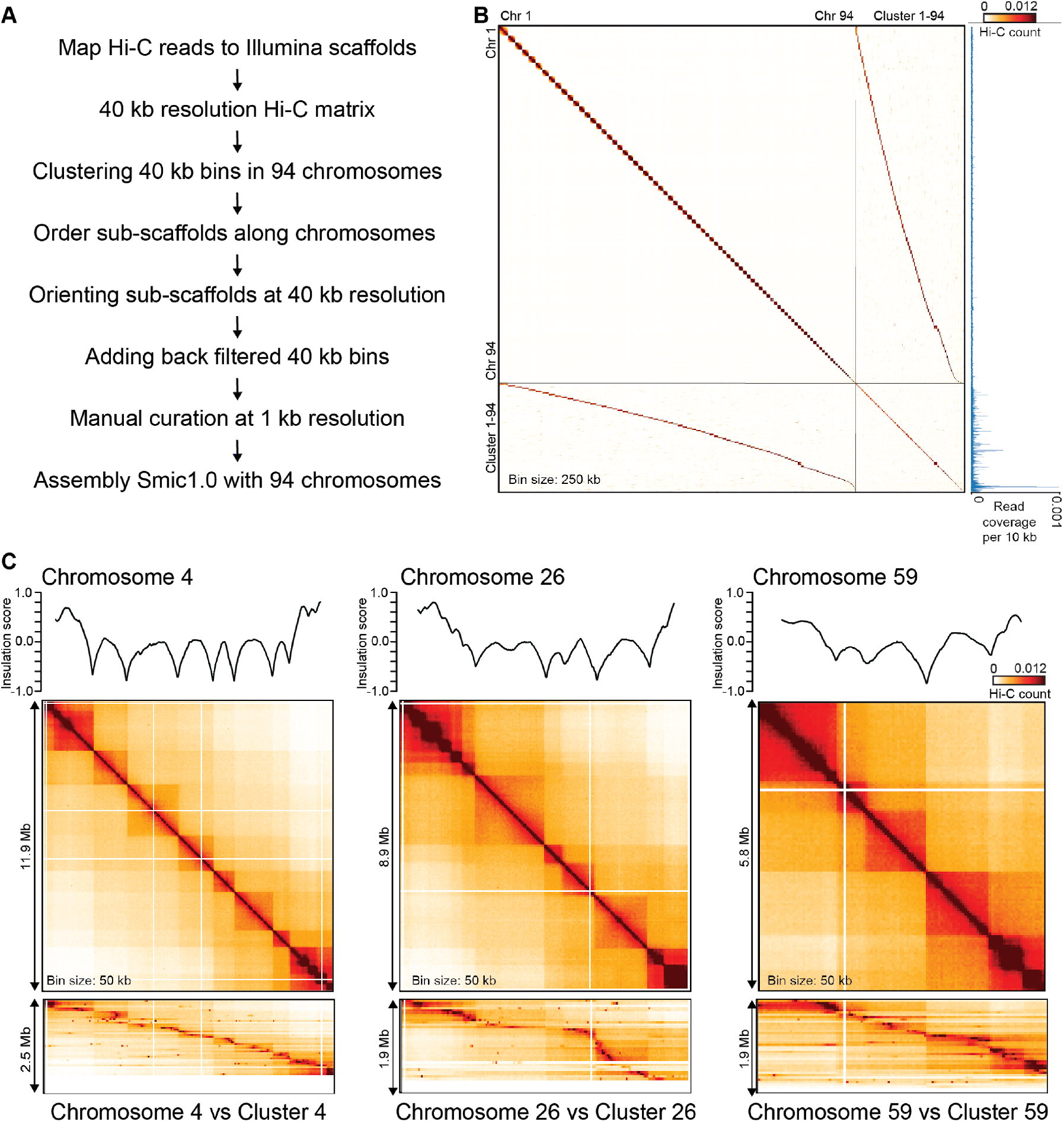
Hi-C assisted assembly of chromosome-scale scaffolds for *S. microadriaticum*. A. Main steps in Hi-C assisted assembly of Smic1.0 B. Hi-C interaction map for the set of 94 chromosomes (ordered by descending size, 250 kb bins) of Smic1.0 and 94 clusters of high copy scaffolds (ordered according to their preferred interactions with the set of 94 chromosomes). Relative sequence coverage per 10 kb bin is shown along the right axis. Each of the clusters interact mostly with only one of the chromosomes, but sequences in these clusters have on average a copy number that is 11 times higher than sequences located along the assembled portions of chromosomes 1-94. C. Examples of Hi-C interactions for chromosomes 4, 26 and 50 (top row of heatmaps). Hi-C data is binned at 50 kb resolution. Plots on top of the heatmaps represent insulation profiles (10 kb resolution, window size 500 kb; see Methods). This profile represents the number of interactions that occur across each location. Local minima in these profiles indicate the locations of sites that strongly prevent interaction to occur across them and these correspond to Hi-C domain boundaries. Bottom row of heatmaps display Hi-C interaction patterns between cluster 4 and chromosome 4, cluster 26 and chromosome 26 and cluster 59 and chromosome 59. Bins in these heatmaps are arranged in order of the position of their most frequent interactions along the corresponding chromosome. Sub-scaffolds present in each cluster may be present as multi-copy arrays, interact mostly with single positions on the corresponding chromosomes, and their interaction patterns follow the domainal patterns seen in the Hi-C interaction maps for each chromosome.

Hi-C reads were mapped to the set of Illumina-based scaffolds (Supplemental Table S1). A total of 2,324,324,062 uniquely mapping chromatin interactions were obtained. The Hi-C data was binned at 40 kb resolution and the interaction matrix was corrected for intrinsic experimental biases ^27^. We used hierarchical clustering (“Karyotyping” ^25^) to identify groups of bins that all interact frequently with each other while interacting much less frequently with other bins. These groups of frequently interacting bins are considered to be located on the same chromosome. After two rounds of hierarchical clustering and manual curation we identified 94 clusters, indicating the presence of 94 chromosomes. Subsequently, we used Hi-C-based “scaffolding” ^25^ to order subscaffolds along each chromosome. Sub-scaffolds are consecutive sections of the original Illumina-based scaffolds that are located on the same chromosome based on Hi-C interaction frequency (see Methods). This approach aims to order sub-scaffolds in such a way that there is an inverse relationship between interaction frequency and genomic distance for all pairs of loci along each chromosome. We then manually corrected the order and orientation of sub-scaffolds along each chromosome (see Methods for details).

After a final manual curation of the assembly using Hi-C data binned at 1 kb resolution, we obtained a genome-wide assembly (termed Smic1.0) that contains 94 chromosomescale scaffolds that combined cover 624,473,910 bp (77% of the starting 808,242,489 bp; scaffold N50 = 8.44 Mb, contig N50 = 23.35 kb; Supplemental Table S2). The chromosome number is close to previous estimates of 97+/-2 chromosomes based on electron microscopic analyses ^28–31^.

### High copy sub-scaffolds

In the process of assembly, we set aside a total of 183,768,579 bp. Many of these sequences were set aside because they interacted frequently with more than one chromosome. Interestingly, we found that these excluded sub-scaffolds could be clustered in 94 groups (referred to here as clusters 1-94) based on their Hi-C interaction frequencies. Accordingly, each of these clusters interacts particularly frequently with only one of the 94 chromosomes, and each chromosome interacts frequently with subscaffolds from only one such cluster (Figure 1B), indicating that these sequences may in fact be part of those chromosomes. Analysis of Hi-C read coverage indicated that while sequences present on the assembled chromosomes 1-94 are all present at similar copy number, many of the sub-scaffolds located in clusters 1-94 were present at much higher copy number (Figure 1B). The higher copy number of these sub-scaffolds can explain in part why they also show relatively high interaction frequencies with other chromosomes. Visual inspection of the Hi-C interaction patterns revealed that each sub-scaffold within a cluster interacts mostly with a single region on the assembled chromosomes (Figure 1C; lower row of interaction maps), indicating that they may be located at or near that single position but often in multiple copies. Consistent with this, in some but not all cases, the high copy sub-scaffolds were part of the same original IIlumina scaffolds as the sub-scaffolds present at the corresponding location of the assembled chromosome. Combined, these analyses imply that some genomic segments are present as multiple copies at distinct positions along each of the 94 chromosomes.

### Hi-C interaction map displays domainal features

The Hi-C interaction maps of all chromosomes show domainal features: each chromosome displays a series of square-shaped domains along the diagonal with relatively elevated interaction frequencies within them and lower frequencies between them. The boundaries between them are often, but not always, sharp transitions. Interactions between loci on either side of a boundary are strongly repressed, and therefore domain boundaries act as structural “insulators” ^32^. Further, interactions between these Hi-C domains reveal an apparent nested series of squares and rectangles farther from the diagonal. Hi-C interactions maps obtained from cultures enriched in coccoid cells (G2/M immobile cells) or mastigote cells (G1/S flagellated cells) revealed no obvious differences (Supplemental Figure S2).

To further analyze these domainal features, we used the previously described chromatin insulation analysis ^32^ to determine the positions of Hi-C domain boundaries genome-wide at 10 kb resolution (Figure 1C). In this analysis loci are identified that strongly prevent interaction to occur across them. We identified 441 boundaries (excluding chromosomes 83-94 that are too short for this analysis). Visual inspection suggests this analysis did not identify some weaker domain boundaries present on most chromosomes. One possible explanation for domain boundaries is the presence of gaps in the Smic1.0 genome assembly. We therefore obtained three independent sets of PacBio long reads ranging in average length from 5.2 to 13.4 kb (Supplemental Table S3) and used these to perform gap-filling using LR_Gapcloser ^33^. The gap-filled genome, referred to Smic1.1, reduced the number of contigs from 44,997 to 10,628 and improved contig N50 from 23 kb to 115 kb (Supplemental Table S2). However, gapfilling also introduced new assembly errors as evidenced by sequences interacting with multiple chromosomes (Supplemental Figure S3). Importantly though, for Smic1.1 we identified the positions of 446 boundaries of which 241 boundaries were located within contiguous sequence (i.e. located at least 15 kb from a contig end). Further, Hi-C interactions of the high copy scaffolds show that these are not predicted to be located at these boundaries (Figure 1). Therefore, we conclude that these boundaries are genuine chromosome structural features (see below). All analyses presented below were performed on the Smic.1.0 assembly and subsequently confirmed on Smic1.1 (Supplemental Figure S4, S5, S6).

### *S. microadriaticum* chromosomes are folded as linearly organized layered rods

Hi-C data can provide insights into the folding of chromosomes. When the average interaction frequency is plotted as a function of genomic distance (*P*(*s*)) a general inverse relationship is typically observed and from the shape and exponent of the curve features of chromosome folding can be inferred. We plotted *P*(*s*) for *S. microadriaticum* (Figure 2A). The shape of *P*(*s*) suggests three regimes. First, for loci separated by a few kb there is a very steep decay. Read orientation analysis shows that the steep decay in regime I is the result of non-informative Hi-C ligation products (^34^; Supplemental Figure S7). Second, for loci separated by several kb up to ~3 Mb there is a very shallow decay (*P*(*s*)~*s* ^−0.4^). Third, for loci separated by more than ~3 Mb there is a sudden steep drop in contact frequency. The overall shape of *P*(*s*) is reminiscent of that observed for mitotic chromosomes in vertebrates, which shows a shallow decay of *P*(s) for loci separated by up to several Mb followed by a steep drop. We have previously demonstrated that such shape of *P*(*s*) is consistent with the formation of a linearly arranged, pseudo-layered organization of relatively stiff rod-shaped chromosomes ^35,36^ where the average size of a pseudo-layer corresponds to the position of the steep drop in *P*(*s*) (Figure 2B). Loci located in different pseudo-layers along the chromosomes only very rarely interact, while loci located within a layer very frequently interact. The Hi-C data suggest that *S. microadriaticum* chromosomes are rod-shaped with pseudo-layers of around 3 Mb: any pair of loci separated by >3 Mb rarely interact, while any pair of loci that are separated by <3 Mb interact frequently. Interestingly, all chromosomes, regardless of their length, show the steep drop in *P*(*s*) at ~3 Mb, which indicates that the layer size is independent of chromosome length.

**Figure 2.**
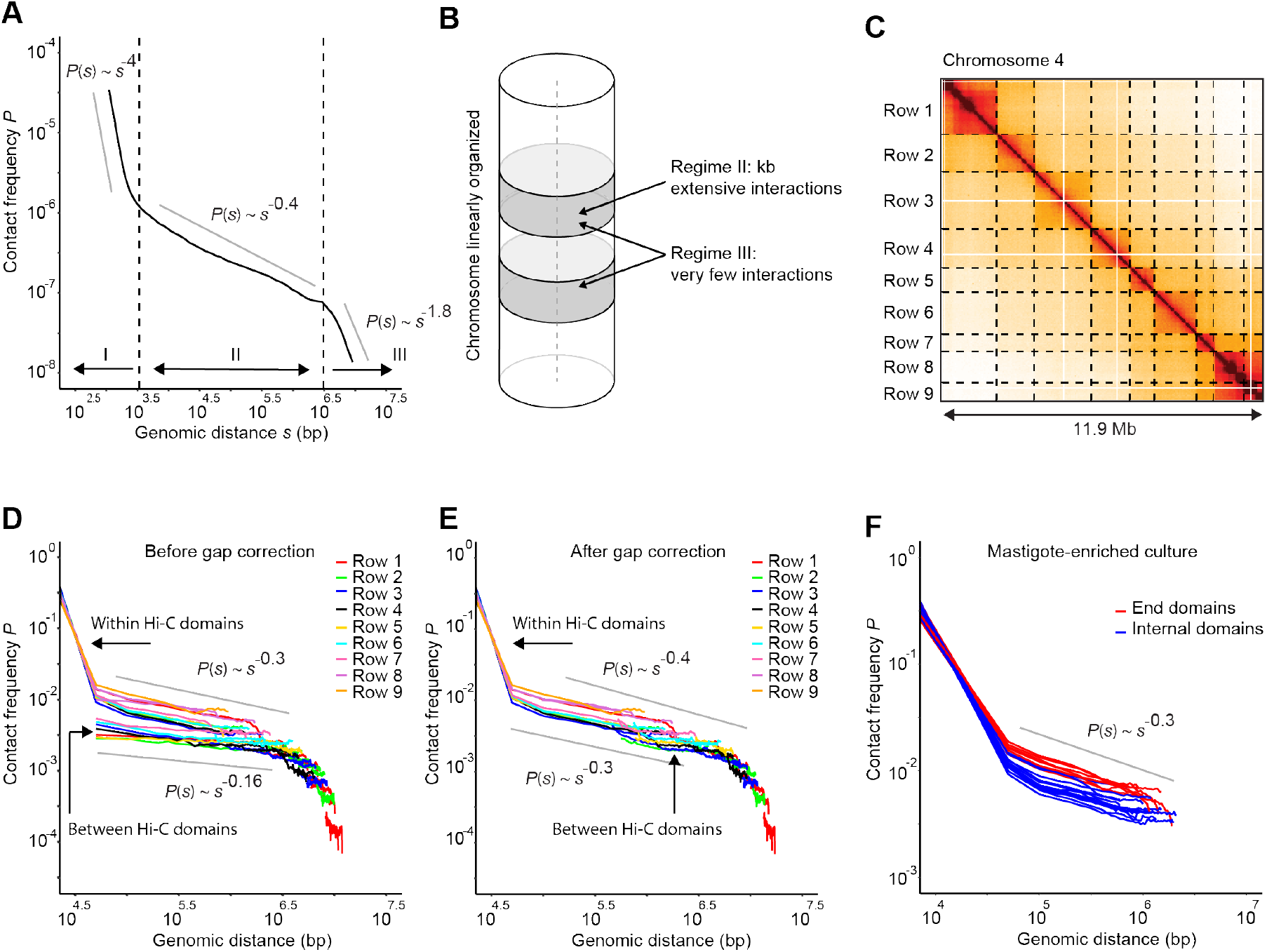
Linear and layered organization of *S. microadriaticum* chromosomes. A. Genome-wide contact frequency *P* versus genomic distance *s* for mastigotes-enriched cultures. *P*(*s*) displays three regimes (I, II and III) with distinct exponents (indicated with gray straight lines). The *P*(*s*) plot is very similar for coccoid-enriched cultures. B. Schematic depiction of a linearly organized and layered chromosome. Linear organization is predicted from the presence of the steep drop in *P*(*s*) (regime III). The shallow decay in interaction frequency followed by a steep drop at ~3 Mb is consistent with a layered organization where layers are around 3 Mb and loci within these layers extensively mix. Regime I is at least in part due to the presence of aberrant Hi-C molecules known to occur at very short distances (unligated ends and self-circularized fragments; see Supplemental Figure S7). C. Hi-C interaction map for chromosome 4 (bin size = 50 kb) for mastigotes. Dotted lines indicate domain boundaries and define a set of squares across the interaction map. D. *P*(*s*) for each square defined by domain boundaries in panel C. Hi-C data from mastigotes. Each individual line represents *P*(*s*) for a single square, colored by row (indicated in panel C). The estimated exponent for *P*(*s*) for regime II ranges from ~−0.16 to −0.3 as indicated by the straight gray lines. Plots for chromatin interaction data within contiguous Hi-C domains, and between Hi-C domains are indicated. E. As panel D, but after correcting genomic distances (*s*) for estimated gap sizes between adjacent Hi-C domains. The estimated exponent for *P*(*s*) for regime II is between −0.3 and −0.4 as indicated by the straight gray line. F. *P*(*s*) plots for Hi-C domains located at the telomeric ends of chromosomes (red lines) and for domains located internally (blue lines) for chromosomes 4, 26 and 59. Hi-C data obtained with cultures enriched in mastigotes. The *P*(*s*) plots are very similar for coccoid-enriched cultures.

The exponent of *P*(*s*) in the intra-layer regime can reveal some properties of how chromatin is organized within each layer. For *S. microadriaticum* the exponent of *P*(*s*) in the intra-layer regime is small: based on the global *P*(*s*) plot shown in Figure 2A the exponent is close to ~0.4. We also plotted *P*(*s*) for interactions that occur within individual Hi-C domains, excluding any interactions that occur across Hi-C domain boundaries (Figure 2C, D). The exponent of *P*(*s*) for interactions within individual domains is consistently ~−0.3 (Figure 2D-E-F-G). Such small exponent indicates extensive packing and potential mixing of DNA within the pseudolayers.

We observe more variable exponents when we plotted *P*(*s*) for interactions between Hi-C domains: exponents ranged from −0.1 to −0.4. When we assume that domain formation is entirely due to sequence gaps, though highly unlikely (see above and below), the *P*(*s*) plots in Figure 2D can be used to estimate the size of such putative gaps by determining how much individual inter-domain *P*(*s*) plots need to be shifted along the x-axis so that they all overlap (see Methods). In most cases plots would need to be shifted several hundred kb. After such putative gap correction the estimated exponents for regime II for the different sections of the Hi-C map again ranged between −0.3 and −0.4 (Figure 2E).

### Fluctuation of chromatin compaction along chromosomes

Visual inspection of Hi-C interaction maps shows that interactions tend to be of higher frequency near the ends of all chromosomes. To quantify this we plotted *P*(s) for telomeric Hi-C domains and for internally located Hi-C domains (Figure 2F). We observed that chromatin interactions within terminal domains are about 2-fold higher for loci separated up to 1 Mb, while the exponent of *P*(*s*) is very similar for all domains (~−0.3). One interpretation is that the chromatin fiber has a shorter contour length near the telomeric ends as compared to chromatin within the middle portions of the chromosomes^37^.

### *S. microadriaticum* has high GC content that increases towards telomeres and decreases at Hi-C domain boundaries

Examining the base composition of the genome of *S. microadriaticum* we observed a remarkably high GC content of 50.51 %, similar to what can be found in some prokaryotes, but certainly much greater than in eukaryotes belonging to the Animalia or Plantae kingdoms ^38^. Interestingly, we find two chromosome-scale patterns of GC content fluctuations, 1) GC content increases towards the ends of the chromosomes (Figures 3A and 3B), and 2) GC content dips to form local minima at Hi-C domain boundaries (Figure 3C). The dip in GC content observed within domain boundaries could suggest that this chromosome architectural feature is encoded within the genome. These findings were also true for the gap-filled genome Smic1.1 (Supplemental Figure S4). Furthermore, considering only Hi-C boundary sequences located within a contig (i.e. at least 15 kb away from the boundaries of a contig) we found the same pattern (Figure 3C, Supplemental Figure S4D and S4E), This provides further evidence that these chromatin domain boundaries and detected by Hi-C and their distinct sequence composition are *bona fide* chromosomal features.

**Figure 3.**
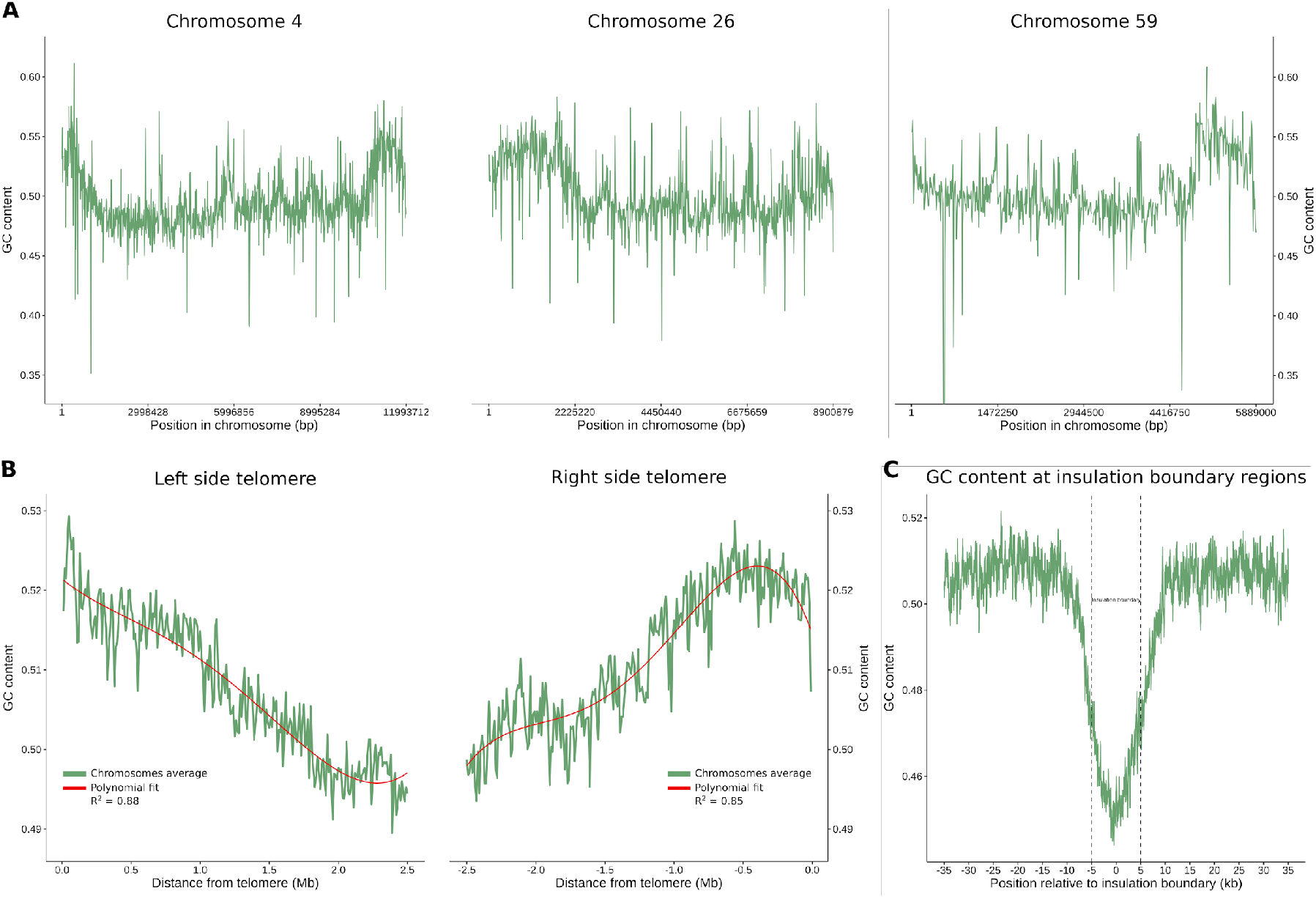
GC content along chromosomes, and near telomeres and Hi-C domain boundaries for Smic1.0. A. GC content fluctuations along chromosomes 4, 26, and 59 measured in 10 kb windows. B. GC content along regions 2.5 Mb from telomeric ends, averaged for chromosomes of sizes of at least 5 Mb and measured in 10 kb windows. GC content decreases as distance to telomeres increases. GC content around insulation boundaries. Values are averaged across all insulation boundaries in the genome for regions 30 kb upstream and downstream insulation boundaries and in 100 bp sliding windows. Dotted lines delimit insulation boundaries. A sharp decline in GC content is observed at insulation boundaries that define Hi-C domains.

### Gene density increases towards telomeres

A chromosome level genome assembly allows to elucidate how genes are placed in a genomic context, which provides insights into higher level organizational principles. This is a particular intriguing point to resolve in dinoflagellates, since their genomes have shown to harbor pervasive gene duplication, some of them arranged in tandem arrays, which may be related to their inertness to transcriptional regulation ^10,14,15,39^. Genome models from Aranda et al ^15^ were mapped to Smic1.0. Of the 49,109 gene models, 48,715, corresponding to 99% of the genes, were successfully mapped using Minimap2 ^40^. When looking at a chromosome level, gene density in *S. microadriaticum* ranges from 38 – 155 genes per Mb, showing a greater gene density compared to other eukaryotic genomes such as human (3.5 – 23 genes per Mb, excluding the Y chromosome) and mouse (7.5 – 15.9 genes per Mb) ^41^. Interestingly, gene density increases towards the telomeres (Figures 4A, 4B and Supplemental Figure S5), having an average gene number of ~ 9 per 100 kb at the end of the chromosomes and decreasing to ~6 towards the central region (Figure 4B). Congruently with gene density, we found that gene expression in *S. microadriaticum* is generally higher towards chromosome ends, resulting in a moderate yet highly significant positive correlation of 0.33 R^2^ and 4.2e^−157^ p-value between gene density and gene expression (Figures 4A, 4H, and Supplemental Figure S5). Furthermore, gene density and expression were positively correlated with GC content (Figure 4H; Supplemental Figure S5). This is in line with the notion that high GC content regions are associated with gene-rich regions.

**Figure 4.**
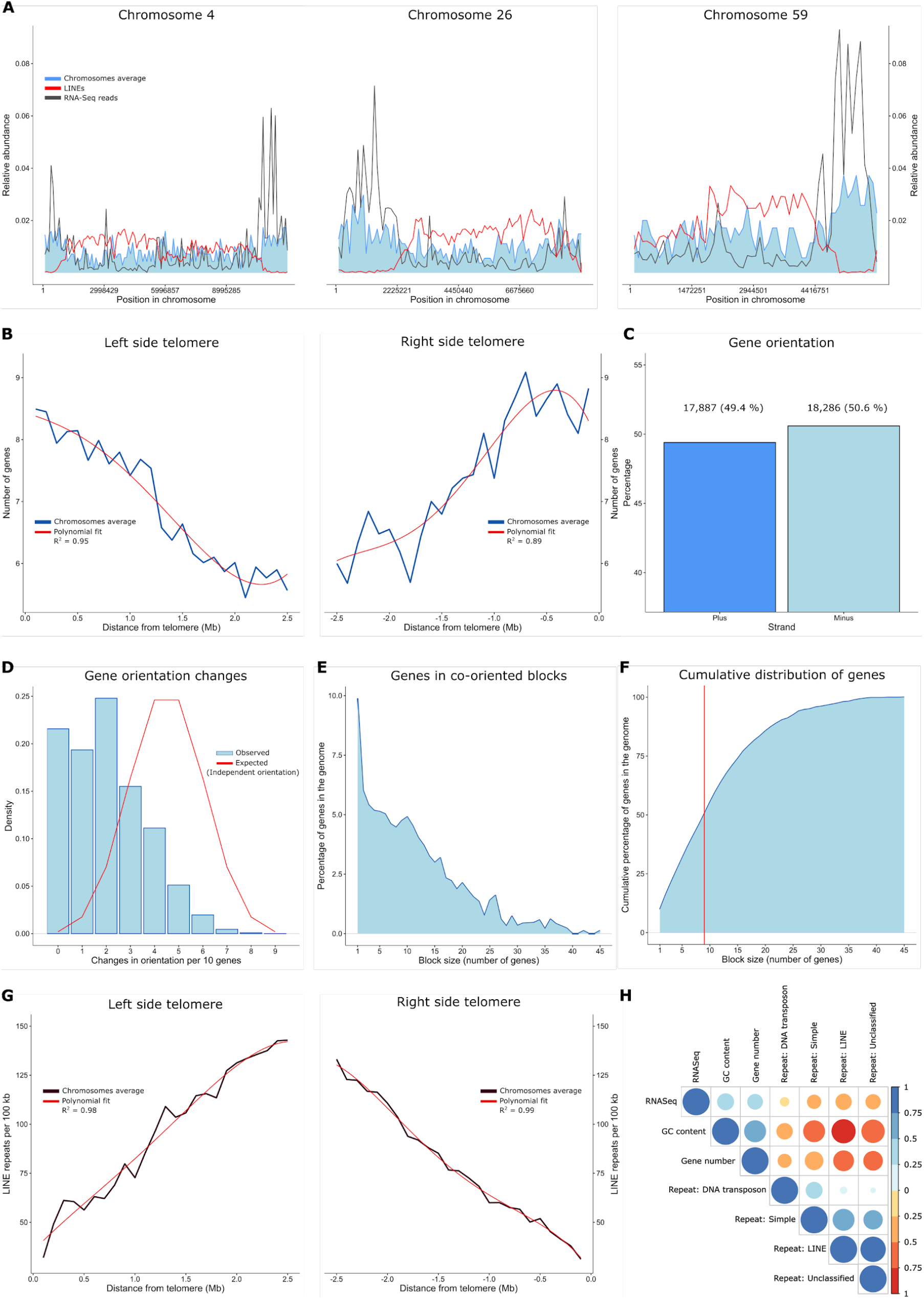
Gene and repetitive element distribution along chromosomes for Smic1.0. A. Relative abundance of genes (blue), LINE repeats (red), and mapped RNASeq reads (grey) for chromosomes 4, 26, and 59. B. Gene number along regions 2.5 Mb from telomeric ends, averaged for chromosomes of sizes of at least 5 Mb and measured in 100 kb windows. Gene number is observed to decrease as distance to telomeres increases. C. Directionality of genes in the genome. A similar number of genes is found in both strands. D. Frequencies of changes in gene orientation. Gene orientation changes defined as the occurrences of neighboring genes located in opposite strands and measured in sliding windows of 10 genes. Observed (blue) and assuming an equal and independent probability of gene orientation (red). E. Distribution of genes in blocks of co-oriented genes. F. Cumulative distribution of genes in blocks of co-oriented genes. 50 % of the genes are found in blocks of 9 or more co-oriented genes (red). G. LINEs number along regions 2.5 Mb from telomeric ends, averaged for chromosomes of sizes of at least 5 Mb and measured in 100 kb windows. LINEs number is observed to increase as distance to telomeres increases. Correlations between Gene number, GC content, RNASeq data, and Repeat types: LINE, DNA transposons, Simple, and Unclassified. Correlation coefficients are displayed as a color and size gradient between positive (blue) and negative (red) values. Correlations were done at 100 kb windows.

### Genes tend to be organized in unidirectional blocks

The organization and orientation of genes along chromosomes shows an even distribution across strands, with 49.4 and 50.6 % of the genes being encoded on the plus and minus strand, respectively (Figure 4C). However, the orientation of neighboring genes is highly correlated and neighboring genes rarely change orientation. Within a 10 gene window, gene orientation changes are strikingly infrequent, similar to our previous analysis ^15^ (Figure 4D). Immediate neighboring genes are more likely to follow the same orientation in prokaryotes, while in eukaryotes orientation of neighboring genes are less or not correlated ^42^. This is largely due to the difference of the polycistronic vs. monocistronic nature of prokaryotic and eukaryotic transcription. In *S. microadriaticum* we observe that genes are preferably organized in blocks of co-oriented genes (Figure 4E), with less than 10 % of the genes found without a co-oriented neighbor. Furthermore, 50% of the genes in the genome are organized in blocks of 9 or more cooriented genes (Figure 4F), showing a high degree of co-orientation at the genome level.

### Chromosomal distribution of repetitive elements

*S. microadriaticum* has a relatively low number of repetitive elements, comprising only 26.5% of the genome compared to 37.5% in mouse ^43^, ~50% in human ^44^, and 84.7% in wheat ^45^. The most abundant repetitive elements in *S. microadriaticum* are long interspersed nuclear elements (LINEs), followed by simple repeats, unclassified repeats, and DNA transposons, constituting 13.36, 5.79, 4.61, and 1.56% of the genome, respectively. Surprisingly, the genome is practically devoid of short interspersed nuclear elements (SINEs) and long terminal repeats (LTRs), together accounting for less than 1% of the genome. We observed that repetitive elements follow the opposite pattern to gene density, expression, and GC-content, being lower towards the ends of the chromosomes and in gene rich regions (Figures 4A, 4G, 4H, and Supplementary Figure S5). The locations of LINE elements are positively correlated with the presence of other repetitive elements, indicating that repetitive elements are in general enriched in the middle part of the chromosomes (Figure 4H).

### Chromatin domain boundaries occur where blocks of unidirectional genes converge

We next investigated the relationship between gene orientation, transcription and features of chromosome conformation observed with Hi-C. Specifically, we were interested to determine whether a relationship exists between transcription of blocks of unidirectional genes and the locations of chromatin domains. Figure 5A shows the Hi-C interaction map for chromosome 19, with domain boundaries indicated by dotted lines.

**Figure 5:**
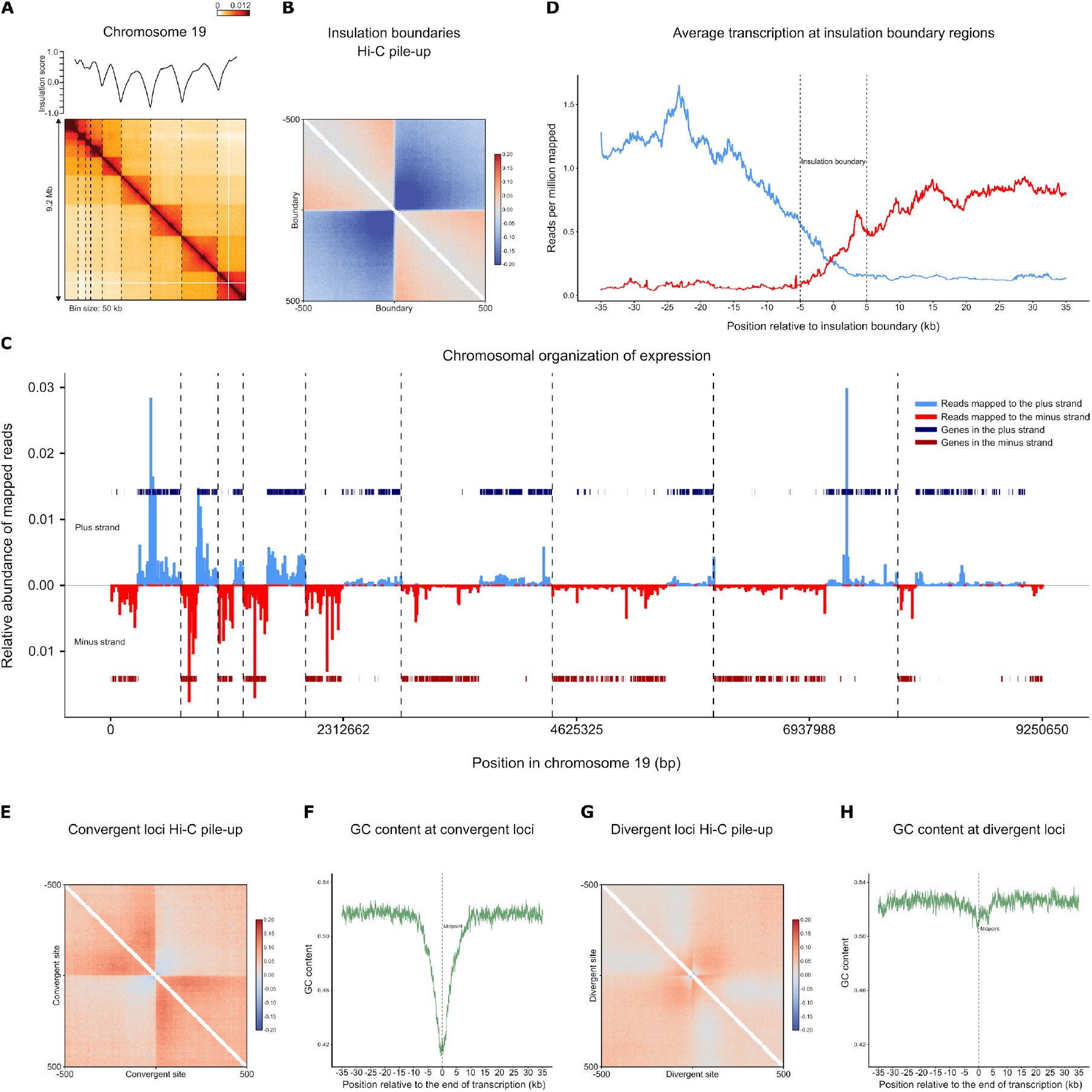
Gene and transcription block organization delimited by domain boundaries. A. Hi-C interaction map for chromosome 19 (bin size = 50 kb) for mastigotes. Dotted lines indicate domain boundaries. Plot on top of the heatmap represent the insulation profile (10 kb resolution, window size 500 kb; see Methods). B. Average Hi-C interactions around the set of 441 domain boundaries at 10kb resolution. C. Chromosome 19 transcription and domain landscape. Indicated are: transcripts mapping to the plus strand (light blue), transcripts mapping to the minus strand (red), genes on the plus strand (dark blue), genes on the minus strand (dark red), and insulation boundaries as dotted vertical lines. A clear domainal gene block organization is observed and is delimited by insulation boundaries. Each domain is a pair of divergent gene blocks. Domain boundaries occur where gene blocks converge. D. Average relative transcription around boundaries. Values are averaged across all insulation boundaries in the genome for regions 30 kb upstream and downstream insulation boundaries and in 100 bp sliding windows. Dotted lines delimit insulation boundary, which is a 10 kb region based on Hi-C data. E. Hi-C pile-up plot for 280 manually curated convergent sites, previously not identified as insulation boundaries, at 10kb resolution. F. GC content around all convergent sites. Values are averaged across all manually curated convergent sites in the genome for regions 35 kb upstream and downstream to the midpoint between two expression blocks (dotted line) in 100 bp sliding windows. A sharp decline in GC content is observed at convergent regions. G. Hi-C pile-up plot for 517 manually curated divergent sites at 10 kb resolution. H. GC content around divergent sites. Values are averaged across all manually curated divergent sites in the genome for regions 35 kb upstream and downstream to the midpoint between two expression blocks (dotted line) in 100 bp sliding windows. Contrary to convergent sites, only a slight decline in GC content is observed in divergent sites.

The sharp boundaries in chromatin interactions are readily detected when we aggregate Hi-C interactions around boundaries genome wide (Figure 5B): interactions across domains boundaries are strongly depleted. To determine whether there is a relation between unidrectional gene blocks and chromosomal domains we plotted RNA expression along each chromosome in a strand specific manner to highlight blocks of co-expressed co-oriented genes (Figure 5C). As expected blocks of transcripts are observed that alternate between being encoded on the top and the bottom strand. Intriguingly, most domain boundaries observed by Hi-C are located at positions where transcription of blocks of unidirectional genes converge. A similar pattern was observed along all chromosomes. To quantify this pattern genome-wide, we plotted the number of reads derived from each strand as a function of distance up or downstream of Hi-C domain boundaries (Figure 5D). We find that reads upstream of a boundary are almost exclusively mapping to the top strand, while reads downstream of a boundary are mostly mapping to the bottom strand.

We did not identify domain boundaries at all locations where transcription of blocks of unidirectional genes converge. This is most likely due to the fact that the parameters we chose for the insulation analysis to identify domains boundaries ^32^ is conservative so that only strong boundaries were reported (see Figure 1, Methods). Visual inspection of Hi-C interaction maps confirms the presence of domain boundaries at most of the locations where gene expression blocks converge. To explore this in another way we identified all sites where transcription of blocks of co-oriented genes converges. This set includes 388 out of the 441 domain boundaries and an additional set of 269 convergent sites that did not overlap a called domain boundary. When we aggregated average Hi-C interactions around this set of additional convergent sites we again observe the formation of distinct structural boundary (Figure 5E). The boundaries are weaker than for called boundaries using the insulation metric (Figure 5B), suggesting these convergent sites are at weaker boundaries that were missed by the stringent domain calling approach (see above). We conclude that domain boundaries occur at the large majority of sites of convergent unidirectional gene blocks, but that some boundaries can be relatively weak. Finally, only in a small minority of cases did we detect a domain boundary away from such convergent sites (53 out of 441 boundaries). These could be due to remaining errors or gaps in the assembly, or they could be different types of structural boundaries.

From these results we conclude that domains are formed by pairs of divergently transcribed blocks of co-oriented genes, with structural boundaries formed where transcription converges. We were interested to explore whether the position within each domain from which transcription of the two blocks of genes diverges displays any particular features. We aggregated Hi-C interactions around the sites from which divergent unidirectional gene blocks are transcribed within each domain (referred to as the bidirectional locus) (Figure 5G). We observe that on average at birectional loci the Hi-C interaction map displays a local insulating boundary: interactions between loci located up to ~100 kb upstream and ~100 kb downstream of the bidirectional locus are depleted. Compared to convergent sites, this boundary effect is much weaker and occurs over only relatively short genomic distances. In addition, we observe lines of enriched interactions that form a “plus’ sign. This can represent long-range looping interactions anchored at the bidirectional locus and other loci located at varying distances either up- or downstream. Whether such loop formation, and positioning of one loop, anchor at divergent loci occurs in *S. microadriaticum* requires more studies. We conclude that both the convergent and divergent sites display specific higher order chromosome structures revealed by Hi-C, with the convergent sites forming very prominent boundaries, and the divergent loci minor and locally acting boundaries. Finally, both types of these display relatively low GC content: a strong reduction is observed at convergent boundaries and a minor reduction is seen at divergent loci (Figure 5F, 5H).

### Some chromosomes are enriched for genes encoding for distinct functional processes

We used GO term enrichment to investigate whether genes located on the same chromosome were functionally related (Supplemental Data Files 4-6). We also checked for tandem arrayed genes with the motivation that genes in such arrays might be linked to related processes. For this reason, we paid particular attention to enriched processes where a majority of available genomic genes were found on any given chromosome. Taking both measures into account, we found genes involved in photosynthesis (chloroplastic), nitrogen-cycling, and stress response (among others) to be enriched on certain chromosomes.

For instance, chromosome 4 contains 7 genes of chloroplastic ATP synthase subunit c genes, 6 of which followed each other in direct vicinity (Smic9977, Smic9979, Smic9980, Smic9981, Smic9983, Smic9984). Furthermore, it contains 16 genes of the pentatricopeptide repeat-containing proteins, organized in 3 clusters, besides additional chloroplastic genes on this chromosome (e.g., PsbP domain-containing protein 7, Short-chain dehydrogenase TIC 32, etc.). Chromosome 62 contained two clusters of Caroteno-chlorophyll a-c-binding protein genes, besides some others encoding chloroplast proteins, and in line with a GO biological process enrichment of photosynthesis (light-harvesting, GO:0009765). Chromosome 69 features a cluster of Caroteno-chlorophyll a-c-binding protein, two clusters of Fucoxanthin-chlorophyll a-c binding protein A, in line with enriched processes related to photosynthesis. Taken together, it seems that chloroplast proteins tend to distribute over few chromosomes, but the functional significance of this is unknown at present.

We also found a surplus of tandem-arrayed nitrogen related genes (12 out of a total of 51) on chromosome 5. Notably, we found several clusters of high affinity nitrate transporters (n = 9 genes, 2 clusters of 6 and 2 genes, and 1 gene) and nitrate reductases (n = 3 genes), some of which were tandem-arrayed, and clusters of Ankyrin repeat domain-containing (n = 6 genes) and Ankyrin-2 proteins (n = 8 genes). Further, chromosome 33 features two clusters of ammonia channel/transporter genes, in line with GO biological process enrichment of ammonium transmembrane transport. Taken together, the data suggest that nitrogen-related genes are arranged in clusters in line with previous speculations that *Symbiodiniaceae* feature extensive gene duplication associated with the provisioning of nitrogen ^15^.

Chromosome 23 is enriched for genes involved in the response to stress, notably signified by a vast expansion of genes annotated as either BTB/POZ and MATH domain-containing protein 2s or BTB and MATH domain-containing protein. For instance, 18 genes encoding for BTB/POZ and MATH domain-containing proteins are concentrated in 1 cluster of 6 genes with the remaining being spread across the chromosome. Further, chromosome 31 contains 40 BTB and MATH domain-containing protein genes in two clusters, suggesting expansion of genes encoding for these proteins in *S. microadriaticum* and revealing their presence in a few specific clusters, as found previously for *Arabidopsis* and rice ^46^. This chromosome further contains 150 genes of chloroplastic pentatricopeptide repeat-containing proteins in various clusters, putatively involved in RNA editing ^47^. At present, it is unclear whether the clustering of functionally related genes along certain chromosomes is a simple consequence of the tandem array arrangement of genes due to mechanisms of duplication or whether this is selected for regulatory reasons.

### High-copy sub-scaffolds are enriched in mobile elements

As shown above, each chromosome contains several high copy sub-scaffolds that appear to be present at multiple copies at specific sites along chromosomes (Figure 1). An overall GO biological process enrichment analysis (i.e., annotating all clusters combined against the genomic background) revealed highly significant overrepresentation of genes associated with DNA integration, reverse transcription, DNA replication, and transposition, suggesting that these sub-scaffolds may represent ‘mobile elements’, which would explain that some appear in high copy number. This was further supported by manual inspection that showed that many of the sub-scaffolds harbored the following genes: ankyrin, copia, retro polymerase proteins/transposons, pentatricopeptide repeat-containing proteins (with exceptions, e.g. cluster 52). While the former three are typical of retrotransposons, pentatricopeptide repeat-containing proteins are commonly found in plants at large numbers (100s in the Arabidopsis and rice genomes), putatively involved in RNA editing ^48^. Besides these overall commonalities, some high copy sub-scaffolds appear to also be functionally enriched for certain genes and their functions not related to transposition. For instance, cluster 1 contains 17 putative surface lipoprotein-encoding genes with similarity to genes in *Shewanella oneidensis*, 10 of which appear in clusters of 6, 2, and 2 genes. Further, cluster 62 harbored 5 genes encoding fucoxanthin-chlorophyll a-c binding protein As (two of which are in an array orientation right next to each other). Six of the said putative surface lipoprotein genes all appeared on a single scaffold of ~ 323 kb in length, corroborating the tandem repeat arrangement of certain genes.

## Discussion

We present a chromosome-scale assembly of the genome of the dinoflagellate *S. microadriaticum*. The genome is composed of 94 chromosomes that range in size from less than 1 Mb up to 16 Mb. This number of chromosomes and their relatively small sizes are in line with microscopic analyses ^28–31^. This assembly reveals the organization of the genetic information and, together with Hi-C data, reveals insights into the spatial organization of chromosomes in this representative of the unique clade of dinoflagellates,

Analysis of the Hi-C interaction maps of *S. microadriaticum* chromosomes reveals several interesting features of chromosome conformation. First, in many eukaryotes Hi-C maps display a distinct plaid, or checkerboard pattern. This pattern reflects the spatial compartmentalization of chromosomes in active and inactive domains ^24^. This pattern arises because active domains spatially interact with other active domains, whereas inactive domains interact with other inactive domains. Hi-C interaction maps for *S. microadriaticum* chromosomes do not display such checkerboard pattern. Thus, compartmentalization of active and inactive regions does not appear to occur in this species.

Second, in many species locus-specific looping interactions are observed. For instance, in mouse and human cells loops can occur between sites bound by the CTCF protein. These loops are thought to be formed through a loop extrusion process mediated by cohesin ^49–54^. Such loops appear as prominent dots in Hi-C interaction maps. Detection of such dots requires deep sequencing of Hi-C libraries (up to 2 billion interactions for the human genome that is ~3 Gb). Our Hi-C interaction maps for *S. microadriaticum* do not reveal dots, despite the fact that we obtained over 2 billion chromatin interactions for this ~0.8 Gb genome. Therefore, it appears that locus-specific loops are not present.

Third, we detect the presence of domainal features: each chromosome is composed of a series of domains that are hundreds of kb up to ~1 Mb in size. Loci located within these domains interact relatively frequently whereas loci located on either side of a domain boundary interact much less frequently. Although some domain boundaries can reflect gaps in the genome assembly, many of these are located within full length contigs indicating these are true chromosome structural features. Furthermore, boundaries display distinct sequence features such as relatively low GC content. We find that each of these domains is composed of a pair of divergent blocks of co-oriented genes and that prominent domain boundaries are located at positions where gene expression converges. This observation suggests a close relationship between gene orientation, gene expression and chromatin domain formation.

Mammalian genomes also display domainal features (Topologically Associating Domains, TADs ^55,56^) that superficially resemble the domains we observe here. However, in mammals TADs do not show a correlation with gene orientation. In yeast small chromosomal domains have been observed that often have boundaries at convergent genes. Both in mammals and in yeast formation of such domains requires the cohesin complex ^52,57–59^. TADs often display looping interactions between their boundaries. The absence of such boundary loops in *S. microadriaticum* suggests that the domains in this species are formed through other mechanisms. The domains we observe here resemble those seen in the prokaryote *Caulobacter* ^60^: they have sharp boundaries, display a similar nested pattern, and no boundary loops. In *Caulobacter* domain boundaries are positioned at highly expressed genes and depend on transcription. Assuming a supercoiled bacterial chromosome, polymer simulations had indicated that domain boundaries can form at sites that block diffusion of supercoils ^60^. However, no relation with gene orientation was reported. Possibly supercoils or plectonemes occur along *S. microadriaticum* chromosomes as well and these may be blocked at sites of convergent gene expression. Future studies are required to test such models or to reveal alternative mechanisms of domain formation in dinoflagellates.

Of note, we find that loci within domains that are located near the ends of the chromosomes display relatively high interaction frequencies, indicating that chromatin folding within those domains differs from domains located more internally. The more frequent interaction within telomeric domains correlates with higher gene density and elevated transcription levels near the chromosome ends. It has been proposed that although the bulk DNA is not wrapped around nucleosomes, actively transcribed genes may have some nucleosomes^19^. If so, the elevated contact frequencies in the gene-rich highly transcribed telomeric domains may be related to the presence of nucleosomes. Alternatively, the telomeric domains may differ in their association with other DNA condensing proteins such as histone-like proteins ^61,62^.

Over the years many models have been proposed on how DNA is spatially organized within dinoflagellate chromosomes. One major hallmark of dinoflagellate chromatin is the observation that most of their DNA is not wrapped around nucleosomes, suggesting the presence of unique chromosome folding mechanisms. Histones are replaced by other basic proteins, e.g. histone-like proteins derived from bacteria and dinoflagellates/viral nucleoproteins (DVNPs) derived from viruses ^20,61,63^.

Microscopically these chromosomes appear as permanently condensed rods, with some variation during the cell cycle ^64^. Chromosomes display characteristic banding patterns. In one model this rod-shaped structure represents a helically coiled toroidal chromonema ^65^. Our data does not support this model: first, the model assumes helical folding, but our Hi-C maps do not reveal such features, which would lead to periodic features in interaction maps, e.g. as seen in prometaphase in chicken cells ^36^. Further, this model assumes circular chromosomes, which is not observed by Hi-C. The optical birefringent properties of dinoflagellate chromatin ^66,67^ suggest that the DNA has liquid crystalline features. This has led to a model where the chromosomes fold as cholesteric liquid crystals ^68–72^. Such crystals form stacks of layers in which molecules are largely oriented in the same direction. In subsequent layers the direction of the orientation of molecules is slightly rotated with respect to each other. Our Hi-C data are consistent with chromosomes forming elongated and relatively stiff rods. We infer this chromosomal shape from analyses where we plot interaction frequency of pairs of loci as a function of genomic distance. In such plots we observe that interactions first decay very slowly (up to 3 Mb) and then drop precipitously for large genomic distances. Previous Hi-C analysis of mitotic chromosomes in mammals and birds and polymer simulations have demonstrated that such biphasic patterns are expected for rod shaped chromosomes ^35,36^.

Loci separated by up to 3 Mb and located within a cross-sectional layer of a chromosome interact frequently and their interaction frequency decays very slowly with increasing genomic distance. The exponent of this decay is around −0.4, and even smaller when *P*(*s*) is analyzed within domains (−0.3), which is much smaller than what is observed in other eukaryotes and even smaller than for mammalian mitotic chromosomes. The protein density on chromatin in dinoflagellates is much lower than in eukaryotes with nucleosome-based chromatin. Therefore, we considered the possibility that the small exponent we observed was the result of inefficient formaldehyde crosslinking of chromatin interactions. Some Hi-C variants use additional crosslinkers such as DSG to increase cross-linking efficiency ^73^. We performed Hi-C using a combination of formaldehyde and DSG. The Hi-C interaction maps and *P*(*s*) plots obtained this way are very similar (Supplemental Figure S8), suggesting the small exponent is not due to low cross-linking efficiency. The small exponent suggests a very high level of compaction but does not by itself reveal how DNA is packed within such layer. Interestingly, the exponent of *P*(*s*) in *Caulobacter* is also remarkably small and similar to what we observe here for *S. microadriaticum*. Possibly plectonemes form within these domains in *S. microadriaticum* as they are proposed to do in *Caulobacter*.

The relationships between the Hi-C domains observed here and microscopically observed structures, such as the banding pattern along liquid-crystalline chromosomes and decondensed loops emanating from the condensed core ^74–76^, remain to be explored. With a chromosome-scale assembly now in hand, locus-specific fluorescent *in situ* hybridization experiments can be designed to probe internal folding of chromosomes and the nucleus in general in more detail.

As to the biological insight obtained from chromosome scale assemblies of dinoflagellate genomes, many of the peculiarities of *Symbiodiniaceae* genomes, such as the unidirectional blocks of genes, the high number of genomic genes, the high density of genes, or the tandem array arrangement of genes (among other) could be corroborated here. This proves that these observations are true at a genome scale level (i.e., across chromosomes). In particular, the observation that a portion of the chromosomes are enriched for specific biological processes (e.g., photosynthesis, nitrogen-cycling, and salt stress) and that functionally related genes tend to co-occur at adjacent sites in the genome add a previously unobserved level of chromosomal functionalization. The enrichment of certain genes along contiguous regions of genomic regions lends support to the hypothesis that two layers of chromosomal ‘specialization’ exist: not only are genes clustered in tandem arrays, but tandem arrays of related genes appear multiple times over a chromosome. It should be noted that deciphering the nature and pervasiveness of this phenomenon is contingent upon a bp-level resolution of the entire chromosomal genomic content, currently unattainable with existing technology and contingent upon the further development of long read sequencing technologies. The comparison of such bp level resolution genomes from different dinoflagellates would then allow to approximate whether the clustering of functionally related genes along chromosomes is an evolutionary selected trait. From an adaptation perspective, and considering the haploid nature of *Symbiodinium microadriaticum* ^12,15^, such a structural organization provides the opportunity for dynamic environmental adaptation through chromosome duplication or loss, pending on how common such alterations are. Varying chromosome counts and polyploidy have been described for field and cultured specimens, in particular autodiploidy during extended culture conditions seems common ^77^.

Despite our inability to completely resolve chromosomes, the remaining 94 ‘clusters’ of sub-scaffolds are likely a mix of sequences of poor assembly and high copy repetitive sequences. However, this does not interfere with our interpretation that tandem gene arrays are likely found in there, but that precise answers as to the distribution, frequency, and architecture of such duplicated regions will have to await bp_resolution of future assemblies (see above). This will hopefully also provide clarity on the relevance of mobile elements and reverse transcriptase on some of these elements, which potentially support the distribution of such clusters. Given that a third (~17,000) of the genomic genes are located in these sub-scaffold regions, a putative very large number of them is duplicated. On the one hand, this might explain the large number of genomic genes in dinoflagellates at large ^15^, on the other hand it may reflect a signature of dinoflagellate adaptation by means of gene/chromosomal duplication/loss. This may in turn constitute part of the answer to the long-standing and broad success of dinoflagellates in the world’s oceans, either as primary producers or as symbiont of marine invertebrates.

## Acknowledgements

We thank Noam Kaplan for advice on Hi-C – assisted genome assembly, Kiruthiga G. Mariappan for initial gene mapping and GO enrichment analysis, and members of the Dekker, Voolstra and Aranda labs for discussions. This work was supported by the Howard Hughes Medical Institute, an endowment by the Byrne family in support of the Joseph J. Byrne Chair in Biomedical Research held by J.D., and by baseline funding from King Abdullah University of Science and Technology (KAUST) to M.A.

## Author contributions

Y.Z. performed all Hi-C and genomic experiments. A.N. performed Hi-C assisted genome assembly and Hi-C analysis. O.R.S performed gene annotation, GC content analysis and RNAseq analysis. M.A, C.R.V. and J.D. supervised the project and contributed to all analyses. All authors contributed to writing of the manuscript.

## Competing Interests statement

The authors declare no competing interests.

## Code availability

cMapping and distiller pipelines for Hi-C analysis: https://github.com/dekkerlab/cMapping https://github.com/mirnylab/distiller-nf

Hi-C-assisted genome assembly: https://github.com/dekkerlab/DNATriangulation

Tools for Hi-C data analysis https://github.com/mirnylab/cooltools

## Data availability

All Hi-C and PacBio sequencing data and Smic1.0 and Smic1.1 genome sequences will be available in GEO. RNAseq data used here are from Liew et al. ^12^ and available in NCBI Short Read Archive (SRA), BioProject PRJNA315758.

## Methods

### *Symbiodinium microadriaticum* culturing

*S. microadriaticum* clade A cultures were obtained from the Gulf of Aqaba near Asia, vendor NCMA (National Center for Marine Algae and Microbiota) – Bigelow Laboratory for Ocean Sciences (CCMP2467-SC) and grown in F/2 FSW (fresh sea water from the gulf of Maine, VENDOR). Four single colonies of *S. microadriaticum* isolated from the original reef sample by growth on F/2 FSW agar plates (clones referred to as D1, D3, D4, D7) were picked and then continued in liquid medium in the presence of a 1:200,000 dilution from a 50X antibiotics stock (100 ml 50X antibiotic stock solution contains 5.0 g Penicillin-G,10.0 g Streptomycin, 5.0 g Kanamycin, 1.0 g Neomycin, 75 mg Nystatin, 30 mg Erythromycin, 40 mg Gentamicin, 80 mg Polymyxin-B, 60 mg Tetracycline, 60 mg Vancomycin). Cultures are grown in T75 tissue culture flasks at 23°C with a 12h/12h light/dark cycle, with a light intensity of 60 - 80 μE m-2 s-1. Once a week cultures were split by first removing the supernatant and adding fresh medium 2.5 hours prior to the start of the light phase. Three hours later (0.5 hours after the light phase has started), the medium with newly born mastigotes is transferred to a new flask and the old vessel is discarded.

### Hi-C procedure

We adapted the conventional Hi-C protocol for analysis of *S. microadriaticum* chromosomes and obtained Hi-C datasets for cultures enriched in mastigotes, and for cultures enriched in coccoid cells (see below). For initial assembly, we pooled all Hi-C data (4 replicates for mastigote-enriched cultures, 4 replicates for coccoid-enriched cultures; Supplemental Table S1) and mapped the reads to the set of scaffolds from Aranda et al. ^15^. Combined, a total of 2,324,324,062 uniquely mapping valid pairs of chromatin interactions were obtained. The Hi-C data was binned at 40 kb resolution and the interaction matrix was corrected for intrinsic experimental biases by balancing using the Iterative Correction method ^27^. Scaffolds smaller than 40 kb were not included in the assembly process.

Below is the Hi-C protocol in detail.

#### Fixation

Hi-C was performed with cultures enriched in mastigotes or enriched in coccoid cells.

##### Mastigotes

Mastigote-enriched cultures were obtained by collecting supernatants of cultures. Cells were fixed with 1% formaldehyde in seawater at RT for 10 min. Fixation was stopped by addition of glycine to a final concentration of 125 mM. Cells were incubated at RT for 5 min, followed by on ice >15 min. Cells were pelleted down and dissolved in seawater. Cell mix was aliquoted into tubes each containing 20 million cells. Cells were pelleted again and incubated on dry ice more than 20 min then stored at −80C.

##### Coccoids

After the removal of the supernatant of cultures (to remove mastigotes), remaining coccoid cells that were attached to the bottom of the flasks were fixed with 1% formaldehyde in 9 ml seawater at RT for 10 min. Fixation was stopped by addition of glycine to final concentration of 125 mM. Cells were incubated at RT for 5 min, followed by on ice >15 min. Cells were pelleted down and dissolved in seawater. Cell mix was aliquoted into tubes each containing 20 million cells. Cells were pelleted again and incubated on dry ice more than 20 min then stored at −80C.

In some experiments Hi-C was performed with cells fixed with formaldehyde and Disuccinimidyl glutarate (DSG). Clone D4 coccoid-enriched cultures were fixed in 1% formaldehyde as described above. After stopping fixation by adding glycine to a final concentration of 125 mM, cells were scraped off the plates. The cells were washed twice in PBS and then resuspended in PBS containing 3 mM DSG. Cells were incubated at room temperature for 40 minutes with rotation. Fixation was stopped by addition of glycine to a final concentration of 400 mM. Cells were incubated for 5 minutes at room temperature, and then pelleted. Cells were washed twice in PBS, pelleted and flash frozen. Fixed cells were then stored at −80°.

#### Cell lysis and restriction digestion

Hi-C was performed on 20 million cells per culture. Cells were resuspended in ~260 μl 1XNEBuffer 3.1 containing protease inhibitors (Thermo Scientific) and then split over 2 Covaris microTubes (Covaris, Part #520045). Cells were then sonicated. For coccoid cells the settings were: 90 seconds, Covaris M220 with the following parameters - peak power 75 watt, duty factor 23 and 200 cycles per burst. For mastigotes the settings were: 20 seconds, Covaris M220 with the following parameters - peak power 75 watt, duty factor 23 and 200 cycles per burst.

Each sample was then transferred to a microfuge tube and 1XNEBuffer 3.1 buffer was added to a total volume of 200 μl. Next, 10 μl 10% SDS was added to a final concentration of 0.5%, and cells were incubated at room temperature for 10 minutes. 23.6 μl 10% Triton X-100 was added to a final concentration of 1%, suspensions were gently mixed and then centrifuged at 3,000 *g* for 5 minutes. The supernatant was discarded, and each pellet was resuspended in 200 μl 1XNEBuffer 3.1. Samples were pooled, centrifuged again and pellet were resuspended in 490 μl 1XNEBuffer 3.1. Each sample was then split over 4 microfuge tubes (118 μl per tube), 8 μl DpnII (400 U, NEB, R0543M) was added to each and samples were incubated at 37°C overnight with rotation.

#### Biotin fill-in of DNA ends, DNA ligation and DNA purification

After overnight digestion, 354 μl 1XNEBuffer 2 was added to each sample. DNA ends were filled in with biotin-14-dATP by adding 60 μl of 1XNEBuffer 3 containing 0.25 mM dCTP, 0.25 mM dGTP, 0.25 mM dTTP, 0.25 mM biotin-14-dATP and 50 U Klenow. Samples were incubated at 23°C for 4 hours in a thermomixer and then placed on ice. DNA was ligated by adding 612 μl of ligation mix (final concentrations in reaction: 50 mM Tris-HCl (pH 7.6), 10 mM MgCl_2_, 1 mM ATP, 1 mM DTT, 5% (w/v) polyethylene glycol-8000, 1% Triton X-10, 0.1mg/ml BSA) and 50 μl T4 DNA ligase (50 units) followed by incubation at 16°C in a thermomixer for overnight. Next, 50 μl 10 mg/ml Proteinase K was added to each sample. Samples were then incubated at 65°C for 4 hours after which 50 μl 10 mg/ml Proteinase K was added again followed by incubation at 65°C overnight. The 4 samples were then pooled in one 15 ml conical tube and mixed with an equal volume phenol-chloroform (1:1). The sample was then transferred to a MaXtract™ tube and then centrifuged at 1,500 *g* for 5 minutes. The aqueous phase was transferred to a clear high-speed Beckman centrifuge tube, and DNA was precipitated by adding 1/10 volume 3M Sodium Acetate, pH 5.2 and 2.5 volumes ice cold 100% ethanol. Samples were incubated at −80°C for at least 60 minutes and then centrifuged at 18,000 *g* at 4°C for 20 minutes. The supernatant was removed, the pellet was dried and then resuspended in 800 μl EB buffer (10 mM Tris-Cl, pH 8.5) and split over two LoBind tubes. DNA was then purified on AMPure beads as follows: 2 volumes of AMPure mix was added followed by 10 minutes incubation at room temperature. Beads were reclaimed using a magnet, supernatant was removed followed by addition of 1 ml 80% ethanol. After 30 seconds of incubation the supernatant was again removed. Beads were washed once more by addition of 1 ml 80% ethanol and then beads were dried at room temperature for 5 minutes. Pellets were resuspended in 50 μl EB buffer and incubated at room temperature for 10 minutes. Beads were reclaimed with a magnet and the supernatant was transferred to a microfuge tube. RNA was removed by addition of 1 μl 10 mg/ml RNase A and incubation at 37°C for 15 minutes.

#### Removal of dangling ends

Biotin was removed from unligated ends by incubating samples (aliquots of 5 μg DNA, typically 10-15 μg per experiment) in 50 μl 1XNEBuffer containing 0.1 mg/ml BSA, 0.025 mM dATP, 0.025 mM dGTP and 15 U T4 DNA polymerase for 4 hours at 4°C. Reactions were then pooled in one LoBind tube, 2 volumes of AMPure mix were added and beads were reclaimed on a magnet. DNA was eluted in 130 μl water.

#### Preparation of Hi-C libraries for Illumina sequencing

DNA samples were transferred to Covaris microTubes, and sonicated for 3 minutes using a Covaris M220 with the following settings: peak power 50 watt, duty factor 20%, 200 cycles per burst. DNA ends were then repaired as follows: 120 μl DNA was mixed with 16 μl 10XNEB ligation buffer, 14 μl 2.5 mM dNTPs, 15 U T4 DNA polymerase, 50 U T4 polynucleotide kinase and 5 U Klenow DNA polymerase in a final volume of 161 μl. Samples were incubated at 20°C for 30 minutes. DNA was then purified by binding to QIAGEN MinElute columns (5 μg DNA per column), washing with 750 μl PE buffer, and DNA was then eluted twice with 17 μl TLE buffer. Next, A-tailing of the DNA molecules was performed by mixing 32 μl of DNA sample with 5 μl 10X NEBuffer 2, 10 μl 1 mM dATP and 15 U Klenow DNA polymerase (3’→5’ exo-). Samples were incubated at 37°C for 30 minutes, followed by incubation at 65°C for 20 minutes. Samples were then placed on ice.

To purify biotin containing DNA fragments, TLE buffer was added DNA samples to make a final volume of 200 μl. Magnetic streptavidin beads (25 μl beads per 5 μg DNA) were washed twice in TWB (Tween Wash Buffer: 5 mM Tris-HCl pH8.0, 0.5 mM EDTA, 1 M NaCl, 0.05% Tween), resuspended in 200 μl 2X Binding Buffer (BB) and added to 200 μl DNA solution. The mixture was incubated at room temperature for 15 minutes with rotation. The beads were reclaimed on a magnet and the supernatant was discarded. Beads were resuspended in 200 μl 1X BB beads and were reclaimed again on a magnet and the supernatant was discarded. Beads were then resuspended in 100 μl 1XT4 DNA ligation buffer (Invitrogen), and transferred to a new tube. The beads were reclaimed on a magnet again, the supernatant was discarded and then resuspended in 40.75 μl 1XT4 DNA ligation buffer (Invitrogen).

To prepare DNA (bound to the streptavidin beads) for Illumina sequencing the DNA sample was mixed with 4 μl Illumina paired end adapters (TruSeq Nano DNA Sample Prep Kit, FC-121-4001), 2.25 μl 5X T4 DNA ligation buffer (Invitrogen) and 3 μl T4 DNA ligase (Invitrogen). All reactions were performed in LoBind tubes. Mixtures were incubated at room temperature for 2 hours. Beads were then reclaimed on a magnet and beads were washed in several steps as follows: first two washes with 300 μl TWB, third wash with 200 μl 1X BB, fourth wash with 200 μl 1X NEBuffer 2 and finally with 50 μl 1X NEBuffer 2. After the last wash the beads were resuspended in 20 μl 1X NEBuffer 2 and then transferred to a new microfuge tube.

DNA was then amplified according to the TruSeq Nano DNA Sample Prep Kit, FC-121-4001 for 6-9 cycles and amplified DNA was purified using AMPure as follows: DNA solution was mixed with 1.1X volume of AMPure XP and incubated at room temperature for 10 minutes. Beads were reclaimed with a magnet and the supernatant was discarded. The beads were then twice washed with 500 μl fresh 80% ethanol. Beads were air-dried for 5 minutes and then resuspended in 30 μl EB and incubated at room temperature for 10 minutes. Beads were reclaimed again and the supernatant was transferred to a new microfuge tube. DNA concentration was then determined by gel analysis.

#### DNA sequencing, Hi-C read mapping and analysis

Hi-C libraries were analyzed by 2X50 bp paired-end sequencing on a HiSeq4000 instrument. Reads were mapped using the cMapping pipeline (https://github.com/dekkerlab/cMapping) or distiller pipeline (https://github.com/mirnylab/distiller-nf). Reads were initially mapped to *S. microadriaticum* scaffolds from Aranda et al. ^15^ to facilitate Hi-C assisted genome assembly, and finally to the assembled genome version Smic1.0.

### PacBio library preparation and sequencing

Genomic DNA was extracted from coccoid-enriched and mastigote-enriched cultures of clone D7 (growing in the presence of antibiotics, see above) using the QIAGEN DNeasy Plant Mini Kits (QIAGEN, Cat# 69104). The cells were ground to a fine powder under liquid nitrogen using a mortar and pestle. Once cell disruption was complete, DNA extraction was performed following the manufacturer’s protocol. DNA from mastigote-enriched cultures was extracted using QIAshredder Mini spin columns while DNA from coccoid enriched cultures was extracted both with and without QIAshredder Mini spin columns.

A first set of DNA libraries were sequencing on a PacBio RS II, and later two libraries were sequenced on a PacBio Sequel I instrument (see Supplemental Table S3). Initial quality control analysis was performed on all samples using Q-Bit, NanoVue, Advanced Analytics-based DNA Fragment Analysis. For initial analysis, material used in libraries analyzed on the PacBio RS II instrument was unsheared as quality control analysis revealed it was already quite fragmented. All samples underwent cleanup steps prior to library construction: Two 0.5X AmpPure bead washes.

In preparation of sequencing DNA on the PacBio Sequel I, DNA was needle sheared: A 1 mL Luer-Lok syringe with 26G 1.5” blunt needles was used for shearing: Sample mD7 (mastigote DNA) was passed 10 passes through the needle, sample cD7 (coccoid DNA), which initially had a smaller starting size and shoulder on the initial quality control, was passed only 5 passes through the needle. Sheared material was assessed using a high-sensitivity DNA Fragment Analyzer assay.

All libraries were of the long-insert genomic DNA type, all constructed using the PB Express 2.0 Kit according to the manufacturer’s instructions. An additional cleanup step was performed following the completion of library construction. Validation quality control analysis performed on all finished libraries included Q-Bit, NanoVue, Advanced Analytics-based DNA Fragment Analysis.

Libraries that were analyzed on the RS II instrument used one SMRTCell with a 10-hour data collection time; libraries analyzed on a Sequel I used one (1M) SMRTCell with a 20-hour data collection time. Read-of Insert (ROI)/ CCS analysis was performed using SMRTLink v.6 or SMRTLInk v.7.

### Genome assembly

We started assembly of the *Symbiodinium microadriaticum* clade A genome with a set of scaffolds generated from Illumina HiSeq reads as described in Aranda et al. ^15^. Combined these scaffolds cover 808,242,489 bp of sequence data over 9,695 scaffolds. The scaffold N50 is 573.5 kb with a contig N50 of 34.9 kb ^15^.

We have generated 4 Hi-C data sets for mastigote - and 4 for coccoid - enriched cultures of *S. microadriaticum* (D1, D3, D4, D7, see above *“Symbiodinium microadriaticum* culturing”) in 2 biological replicates. The 16 datasets were pooled, yielding a total of 4,940,728,852 reads that were then used for Hi-C-assisted genome assembly. A schematic outline of the process of Hi-C-assisted genome assembly is shown in Supplemental Figure S1, and described in detail below. Hi-C data was mapped to the set of scaffolds using the standard cMapping pipeline [^34^ and https://github.com/dekkerlab/cMapping]. Out of a total of 4,940,728,852 Hi-C paired-end reads, for 2,324,324,062, i.e. 47.04%, both ends uniquely mapped to scaffolds. This is comparable to Hi-C data for the human genome where the fraction of uniquely mapping paired end reads is typically around 60%, indicating that the set of scaffolds represents a large majority of the *S. microadriaticum* genome. Of the set of uniquely mapped paired-end reads 1,379,534,687 (59.35%) represented interactions between scaffolds and 944,789,375 (40.65%) represented interactions within scaffolds. Hi-C data was then binned at 40 kb resolution. The final assembly Smic1.0 has 94 chromosomes covering 624,473,910 bp. In addition, for each chromosome we identified a set of sub-scaffolds that are present as high copy number sequences which made correct positioning of them along the chromosomes difficult. Combined these high copy number sub-scaffolds cover 183,768,579 bp.

#### Removal of small scaffolds and bins at ends of scaffolds that are smaller than 40 Kb

Scaffolds smaller than 40 kb were removed due to the relatively low read coverage in Hi-C datasets. Low read coverage will affect normalizing the Hi-C interaction matrix after balancing. Out of 9,695 scaffolds, 7,671 scaffolds smaller than 40 Kb were removed. After binning Hi-C data at 40 kb resolution, each scaffold larger than 40 kb will have a last bin at their 3’ end that is smaller than 40 kb. Those so-called “hanging bins” were also removed. Combined 9,695 bins were removed covering ~70 Mb. The remaining interaction matrix contained 18,468×18,468 bins of 40 kb covering 738,720,000 Mb. The Hi-C interaction matrix was then normalized for technical biases by balancing using the conventional Iterative Correction and Eigenvector decomposition (ICE) method ^27^.

#### Karyotyping

Loci (bins) interact more frequently with other loci located along the same chromosome (in cis) than with loci located on other chromosomes (in trans). This feature can be leveraged to identify sets of bins that all interact with each other more frequently than with others and thus are present on the same chromosome. We refer to this step as karyotyping and it involves converting the Hi-C interaction matrix into a genomic distance matrix followed by bootstrapped clustering of bins based on the distances between pairs of 40 kb bins. We used the algorithm and code as described in Kaplan et al ^25^. We run the clustering 100 times randomly picking 90% of the data in each iteration and then estimated the number of clusters by identifying the largest average distance step in the hierarchical trees which occurred at 82 clusters. We assume that each of these clusters represents a set of loci (40 kb bins) located on the same chromosome.

#### Removal of erroneous bins

We noticed the presence of 40 kb bins that clustered with a set of other bins but also displayed high interactions with bins within other clusters, indicating they contained sequences present on two different chromosomes. These bins may contain “misjoins” where the original scaffolds contained incorrectly joined sequences from two chromosomes. Bins were assumed to be erroneous when the sum of their interactions with bins outside their cluster was above 300 counts. These bins were removed. This led to the removal of 5,239 bins covering 209,560,000 bp. The remaining interaction matrix consisted of 13,229×13,229 bins of 40 kb covering 529,160,000 bp.

#### Creation of sub-scaffolds

Bins that were clustered together, did not contain misjoins and that were adjacent within the same original scaffold were merged to form “sub-scaffolds”. Sub-scaffolds are parts of original scaffolds where they were linked to other sub-scaffolds by misjoins. Subscaffolds are high confidence scaffolds: they were originally assembled using short reads by Aranda et al., their 40 kb bins were found to be located on the same chromosome (cluster) by Hi-C, and they do not contain misjoins using the threshold described above. We created 3,202 sub-scaffolds again performed karyotyping to group sub-scaffolds located along the same chromosome and again identified 88 clusters. 10 subs-scaffolds (400 kb of sequence) were removed from the matrix at this step by DNA triangulation filters. Hence we assembled 3,192 sub-scaffolds that combined cover 528,760,000 bp.

#### De novo scaffolding

Next we set out to order sub-scaffolds along chromosomes. To this end we mapped the pooled Hi-C data to individual sub-scaffolds and binned the Hi-C data so that each bin contains a full length single sub-scaffold. The interaction map was then balanced using ICE ^27^ to create a normalized interaction matrix of 3,192×3,192 bins. We then used Hi-C interaction frequencies between sub-scaffolds within each cluster to order them along chromosomes in a process referred to as “scaffolding” as described by Kaplan ^25^. Scaffolding is based on the fact that Hi-C interaction frequencies decay with genomic distance. We modified the previously published scaffolding algorithm ^25^ and used a probabilistic model that assumes that the distance dependent decay follows a power law to find sets of likely sub-scaffold positions for each chromosome. The solution space is not concave and individual solutions may represent local minima. Therefore, we repeated the optimization with 10 different starting points and through 1000 iterations with an optimization algorithm L-BFGS (Limited-memory Broyden–Fletcher–Goldfarb–Shanno algorithm) we identified the best solution which was reported as the ordering of sub-scaffolds within each cluster. Next we created a new interaction matrix where each chromosome was composed of the newly ordered sub-scaffolds split in their original 40 kb bins. The new chromatin interaction matrix has 13,219×13,219 bins of 40 kb = 528,760,000 bp covering 88 chromosomes (Matrix expansion step).

#### Manual correction of clustering, ordering and orientation of sub-scaffolds

Visual inspection of the chromatin interaction matrix revealed errors. First, some 40 kb bins were assigned to the incorrect cluster. We manually assigned such bins to the cluster it interacts most strongly with. Second, errors in ordering of sub-scaffolds along chromosomes were visible by sharp breaks and checkered appearance of the interaction maps characteristic for inversions and re-arrangements ^78^. These errors were manually corrected to create chromatin interaction maps with smooth distance dependent decay in Hi-C interactions. Third, all sub-scaffolds were oriented according to the original scaffold sequences and thus about half were expected to be in the incorrect orientation. For sub-scaffolds represented by multiple 40 kb bins the orientation could be inferred by examining interactions with their flanking sub-scaffolds. We manually oriented each of the >40 kb sub-scaffolds so that their interactions with flanking sub-scaffolds followed a smooth distance dependent decay. Sub-scaffolds composed of a single 40 kb bin are oriented below.

For all manual corrections of the assembly, at this stage and below, we attempted to maintain the continuity of bins within sub-scaffolds and only moved bins away from other bins of the same sub-scaffold when the Hi-C interaction pattern was very obviously incorrect.

#### Adding back previously removed bins

In the beginning of the assembly process we had removed 40 kb bins that displayed interaction counts >300 with clusters to which they were not assigned (“erroneous bins”, above). We noted that some of these bins were consecutive “chunks” of 80 or more kb within the original scaffolds. We reasoned that many of such large chunks of multiple 40 kb bins would not contain misjoins and could be placed in the current assembly. To this end we created a new fasta file composed of the 88 clusters assembled above, as well as the sequence of all erroneous chunks of 80 kb or larger (479 chunks). We mapped the Hi-C data to this sequence and balanced the interaction matrix using ICE ^27^. We then applied the karyotyping procedure to the 2,543×2,543 40 kb bin interaction matrix containing the chunks and identified 77 clusters. Each of these 77 clusters was then manually assigned to one the 88 clusters assembled above, and inserted in their appropriate locations and orientation to accommodate smooth distance dependent decay of their interactions with the other sub-scaffolds present. In addition, we identified 3 clusters of chunks that did not interact frequently with any of the 88 clusters assembled above indicating that these represent 3 additional chromosomes. We added these three clusters as separate chromosomes so that the total number of clusters / chromosomes at this stage of the assembly was 91. In total we added 2,543 bins of 40 kb (101,720,000 bp) back into the assembly that now covers 630,480,000 bp. We also were able to place 4 bins of 40 kb that had been removed by triangulation filters above so that the assembly totals 630,640,000 bp.

After adding back the erroneous chunks we re-evaluated the cluster assignment of each bin. We identified 180 bins of 40 kb that were clearly mis-assigned and manually placed them within the correct cluster they interacted most frequently with.

#### Orienting sub-scaffolds composed of single 40 kb bins

The assembly contains 374 40 kb bins that either represent a single full sub-scaffold, or that had been manually separated from other 40 kb bins from the same sub-scaffold to ensure smooth distance dependent decay in Hi-C interactions. In order to orient these singletons we first mapped the Hi-C data to the current assembly and binned the data at 8 kb resolution so that interactions between the left end and the right end of these 40 kb sub-scaffold with flanking sequences could be manually evaluated and the subscaffolds could be oriented to ensure smooth distance dependent decay.

#### Adding back hanging bins

As outlined above during the first steps of the assembly process the final 3’ end bins of the original scaffolds from Aranda et al. ^15^. that were less than 40 kb (“hanging bins) were removed. In total 9,695 hanging bins covering 69,522,489 bp were left out of the assembly. At this step these hanging bins could be placed back in the appropriate position and orientation if the adjacent bin from the original scaffold was present in the assembly. In total 1,541 hanging bins covering 30,548,811 bp could be placed back into the assembly that now makes up 661,188,811 bp. A new fasta files was created, and the pooled Hi-C data was mapped to this genome and interaction data was now binned at 1 kb resolution.

#### Manual curation of genome assembly at 1 kb resolution

In a final refinement of the assembly we assessed Hi-C interaction frequencies at 1 kb resolution. We removed 1 kb bins when the sum of all their interaction frequencies with other bins along the same chromosome was below 60% of the most frequent (Max) sum of all intra-chromosomal interactions for all 1 kb bins along the same chromosome (see figure below). Such 1 kb bins display relatively high interaction frequencies with one or more other chromosomes and may contain sequences that are incorrectly included in the scaffolds generated by Aranda et al. ^15^

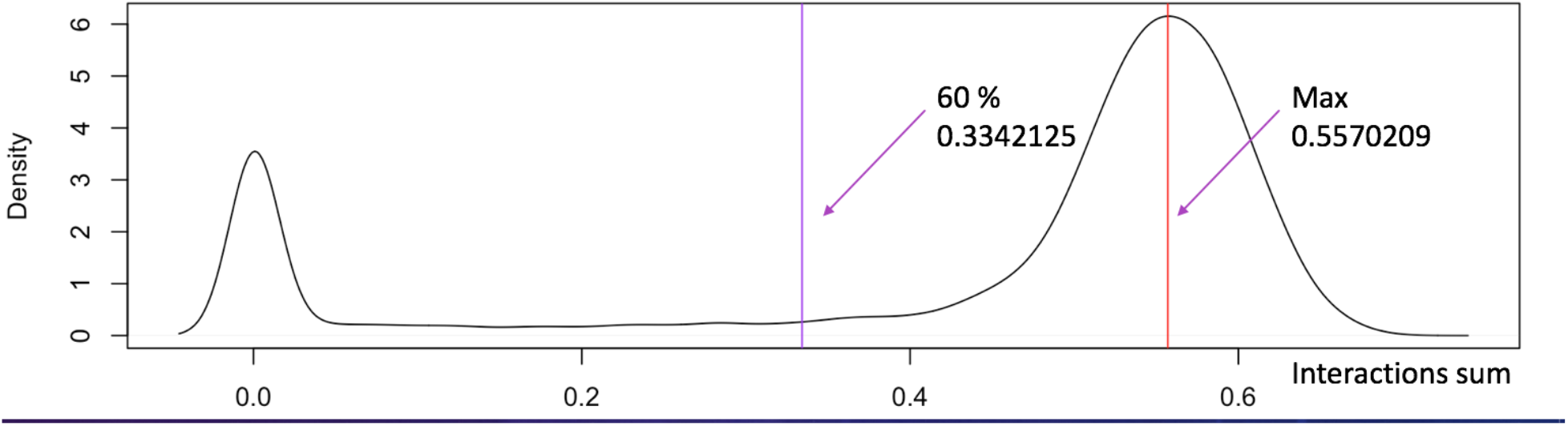

At this step 32,400,317 bp were removed from the assembly. In addition, two clusters were deemed to be composed of 2 and 3 clusters respectively and split accordingly, adding 3 new clusters (chromosomes) to the assembly. At this stage the assembly contains 94 chromosomes covering 628,788,494 bp.

#### Final manual removal of erroneous bins & correction of bin orientations

The last step of the assembly involved visual inspection of Hi-C interaction maps of all 94 chromosomes at 1 kb resolution. Any 1 kb bin for which the Hi-C distance dependent decay pattern appeared erroneous was removed. In total 4,314,584 bp were removed.

#### Final assembly Smic1.0

The final assembly contains 94 chromosomes covering 624,473,910 bp. We arranged chromosomes in order of decreasing size of the clusters to obtain assembly version Smic1.0.

#### Cluster 95: Non-assembled high-copy sequences

All sequences from the original set of scaffolds assembled by Aranda et al. ^15^ that could not be placed on chromosomes 1-94 (in total 183,768,579 bp) were concatenated to form “cluster 95”.

#### Copy number analysis

Hi-C raw single end read coverage analysis of chromosomes 1-94 and cluster 95 revealed that loci along the 94 assembled chromosomes displayed very similar copy number. In contrast, sequences in cluster 95 displayed higher copy numbers (average 11X higher than sequences on chromosomes 1-94). This indicates that the (sub-) scaffolds that make up cluster 95 are present at high copy number and this may explain in part the fact that these could not be placed consistently or with confidence at defined positions along chromosome 1-94.

#### Assigning high copy sub-scaffolds to chromosomes 1-94

To further investigate sub-scaffolds that make up cluster 95, we re-mapped the Hi-C data using the distiller pipeline (https://github.com/mirnylab/distiller-nf) to a new fasta file which includes chromosomes 1-94 and all remaining sub-scaffolds which were left out of the assembly as separate entries to the fasta file (31,552 sub-scaffolds). We then binned and Iced the data at 1Mb resolution. Note that because sub-scaffolds are much smaller than 1 Mb, each will simply contain a single full length sub-scaffold. Next, we calculated the average size-normalized Hi-C interaction frequency between each bin and each of chromosomes 1 through 94 to identify the chromosome each sub-scaffold bin interacts with mostly. In several cases we could not assign the chromosome a given sub-scaffold interacts mostly with (e.g. due to zeros in the contact map). In those cases we assigned such sub-scaffolds to the cluster to which the previous sub-scaffold from the same original Illumina-based scaffold was assigned. Sub-scaffolds that all interact with the same chromosome were then concatenated to form a single set. Sub-scaffolds that could not be assigned to any specific chromosomes were concatenated to form a separate set (number 95). Finally, a new fasta file was created that contained chromosomes 1-94, followed by 94 sets of sub-scaffolds that interact with chromosomes 1-94 respectively, followed by a final set (number 95) of sub-scaffolds that could not be assigned to any chromosome. This fasta file includes all the sequence i.e. 808,242,489 bp that made up the set of Illumina-based scaffolds generated by Aranda et al. ^15^.

#### Smic1.1: filled gaps in Smic1.0 using PacBio long read data

We sequenced *S. microadriaticum* genomic DNA (coccoid clone 7 and mastigote clone 7) on 8 SMRT cells of a RSII PacBio sequencer. We also sequenced genomic DNA of each prepartion on 1 SMRT each on a PacBio Sequel II. In total out of these 10 SMRT cells we obtained 3,732,095 subreads out with an average length of ranging from 3.4 kb to 13 kb (Supplemental Table S3). We have used Minimap2 ^40^ and BLASR ^79^ for mapping these PacBio subreads to Smci1.0 using default settings. We have found 3,239,652 unique mapping reads with Minimap2 and 3,359,291 unique mapping reads with BLASR out of 3,732,095 reads (Supplemental Table S3).

To fill gaps in the Smic1.0 genome, we used the 3,732,095 PacBio subreads and LR_Gapcloser ^33^ running for 6 iterations and under default settings. The number of Ns in the assembly was reduced from 7.378 to 0.396 % of the assembly, scaffold number decreased from 44,997 to 10,628, and the contig N50 increased from 23.35 to 115.858 kb (Supplemental Table S2). The gap-filled genome assembly is referred to as Smic1.1.

### Hi-C domain boundary detection

For the scaling plot for Hi-C domains, the positions of Hi-C domains were first defined by their boundaries using matrix2insulation script from cWorld (https://github.com/dekkerlab/cworld-dekker/blob/master/scripts/perl/matrix2insulation.pl) using all combined data matrix file binned at 10 kb resolution ^32^. The insulation window size was 500 kb. This analysis produces an insulation profile along chromosomes (examples of insulation plots are shown in Figure 1C). Local minima in insulation profiles indicate the positions of Hi-C domain boundaries, and the script produces a list of such boundaries and their strength ^32,34^. To define a set of high confidence Hi-C domain boundaries we selected boundaries with a boundary strength equal or greater than 0.2. Given that local minimum detection has an error of around +/- 1 bin, we manually corrected all boundaries calls (i.e. shifting the positions of boundaries 1-2 bins) based on visual inspection of the Hi-C interaction map. The final list included 441 Hi-C domain boundaries (not including chromosomes 83-94 which are too small for insulation analysis with the settings described above). For Smic1.1 total 446 boundaries were found (not including chromosomes 83-94).

To check boundaries within contigs (less than 24 consecutive N’s in a 30 kb window around the boundary), from the above list: boundaries within subscaffolds were filtered and then filtered again with 10kb upstream and 10kb downstream adding to the 10 kb boundary (30 kb total) where this 30 kb region has less than 24 consecutive N’s. For Smic1.0 we have 65 such boundaries and for Smic1.1 we identified 241 boundaries located within contigs.

### *P*(*s*) calculations and estimation of gap sizes between Hi-C domains

*P*(*s*) plots were calculated in two ways. First, *P*(*s*) was calculated genome-wide using valid chromatin interaction pairs with the following script from cooltools package with all default settings (https://github.com/mirnylab/cooltools).

For *P*(*s*) calculations for single chromosomes at the level of chromosomal Hi-C domains we used Hi-C data binned and balanced at 50 kb resolution. Hi-C domains borders were calculated by insulation analysis (see above). The grid of domain borders define a set of squares throughout the Hi-C interaction map. *P*(*s*) was calculated for each square by plotting the average of each diagonal of 50 kb bins within the square as a function of *s*. For all squares not centered at the main diagonal, the values of *P*(*s*) for the smallest and largest *s* were left out because they are calculated only for 1 bin and thus noisy.

When we assume that Hi-C domains are the result of gaps in the assembly, we can estimate the size of the gaps by calculating what the genomic distance should be between two loci immediately adjacent of a Hi-C domain boundary given the observed interaction frequency between them. To estimate the sizes of gaps for a few chromosomes we calculated *P*(*s*) plots for all on-diagonal Hi-C domains for those chromosomes and for the squares in the Hi-C interaction maps that are positioned immediately off-diagonal and represent interactions between adjacent Hi-C domains. We then estimated the sizes of the gaps at the boundaries between Hi-C domains by determining how much the *P*(*s*) plots of the off-diagonal squares needed to be shifted along the x-axis to make them smoothly overlap with the *P*(*s*) plots of the on-diagonal domains. Interestingly, application of gaps estimated in this manner make *P*(*s*) plots of all squares of the Hi-C maps overlap more smoothly. This included *P*(*s*) plots for squares of the Hi-C maps that correspond to interactions between Hi-C domains that are separated by more than 1 boundary/gap.

### Genome annotation and analysis

#### Identification and masking of repetitive elements

Repetitive elements in the Hi-C scaffolded genome were identified and masked with RepeatMasker (Smit A, Hubley R, Green P. *RepeatMasker Open-4.0* 2013-2015 [Available from: http://www.repeatmasker.org) using the *de novo* repeat library for *S. microadriaticum* generated by Aranda et al. ^15^. This resulted in masking 26.45 % of the genome, of which the most abundant repetitive elements were LINES, DNA transposons, simple repeats, and unclassified. More than 50 % of the repetitive elements were LINEs, comprising 13.36 % of the genome. To observe the distribution of repetitive elements along chromosomes, we measured the abundance of the most prominent repetitive elements using 100 kb non-overlapping windows.

#### Genome annotation, enrichment analyses, and gene expression

As the Hi-C scaffolded genome is based on the previous *S. microadriaticum* assembly ^15^, the annotation of the Hi-C scaffolded genome consisted of remapping the annotation of the previously generated genome by Aranda et al. ^15^ to the Hi-C scaffolded genome with Minimap2 ^40^. 48,715 out of 49,109 genes were mapped from the original assembly to the new Hi-C scaffolded genome. GO enrichment analysis was done at a chromosome level with topGO (version 2.37.0; Bioconductor package Alexa A, Rahnenfuhrer J (2020). *topGO: Enrichment Analysis for Gene Ontology.*) and evaluated with weight01 Fisher statistic at a 0.05 p-value threshold. Z-scores were calculated for each GO term per chromosome using inhouse scripts. KEGG orthology was assigned with BlastKOALA (28), mapped to functional pathways using GAEV ^80^, and tested for pathway enrichment using a hypergeometrical distribution and corrected for multiple testing with Benjamini-Hochberg procedure at a 0.05 p-value threshold. To assess gene expression across different regions of the chromosomes, RNASeq reads previously generated by Aranda et al ^15^, were mapped to the Hi-C scaffolded genome using HiSAT2 (v. 2.1.0) ^81^.

#### Gene distribution and orientation analyses

Gene distribution along chromosomes was measured as the number of genes found in 100 kb non-overlapping windows. The distribution of genes in blocks of co-oriented genes was measured by counting the number of consecutive genes found on the same strand (plus or minus) until the next neighboring gene appeared on the opposite strand. Gene orientation changes were measured as the number of times neighboring genes appeared on opposite strands within a 10 gene sliding window and compared to orientation changes assuming an equal probability and independent occurrence of genes at either strand using a binomial distribution.

#### GC content

GC content was measured only in defined regions where at least 50 % of the bases were [A,C,G,T], meaning that regions containing 50 % Ns were not used for the analysis. GC content along chromosomes was measured in 10 (for plotting) and 100 (for correlation analysis) kb non overlapping windows. GC content surrounding insulation boundaries was measured in 100 bp sliding windows across 70 kb regions that included 30 kb upstream and 30 kb downstream of the 10 kb insulation boundaries.

#### Telomeres analyses

Analyses of telomeric regions were done using 2.5 Mb from each telomeric end, meaning that chromosomes smaller than 5 Mb were not included in the analyses. This resulted in 69 chromosomes utilized for the analyses. Gene number, gene directionality, and LINEs number were measured at window sizes of 100 kb, whereas GC content was measured at 10 kb windows. For every plot of each analysis, a polynomial of the fourth order fit was derived together with the respective coefficient of determination (R^2^).

#### Correlations at a chromosome level

Correlations between Gene number, GC content, RNASeq, and repetitive elements: LINEs, DNA transposons, Simple repeats, and Unclassified repeats, were performed using values from 100 kb non-overlapping windows. Correlations were performed in R with the corrplot package (v. 0.84) using a Pearson’s correlation and corrected for multiple testing with Benjamini-Hochberg procedure at a 0.05 p-value threshold.

## Notes

### Competing Interest Statement

The authors have declared no competing interest.

