## Supplemental Materials for "Chromosome-scale assembly of the coral endosymbiont *Symbiodinium microadriaticum* genome provides insight into the unique biology of dinoflagellate chromosomes"

**SUPPLEMENTAL FIGURES**

**SUPPLEMENTAL FIGURE 1**

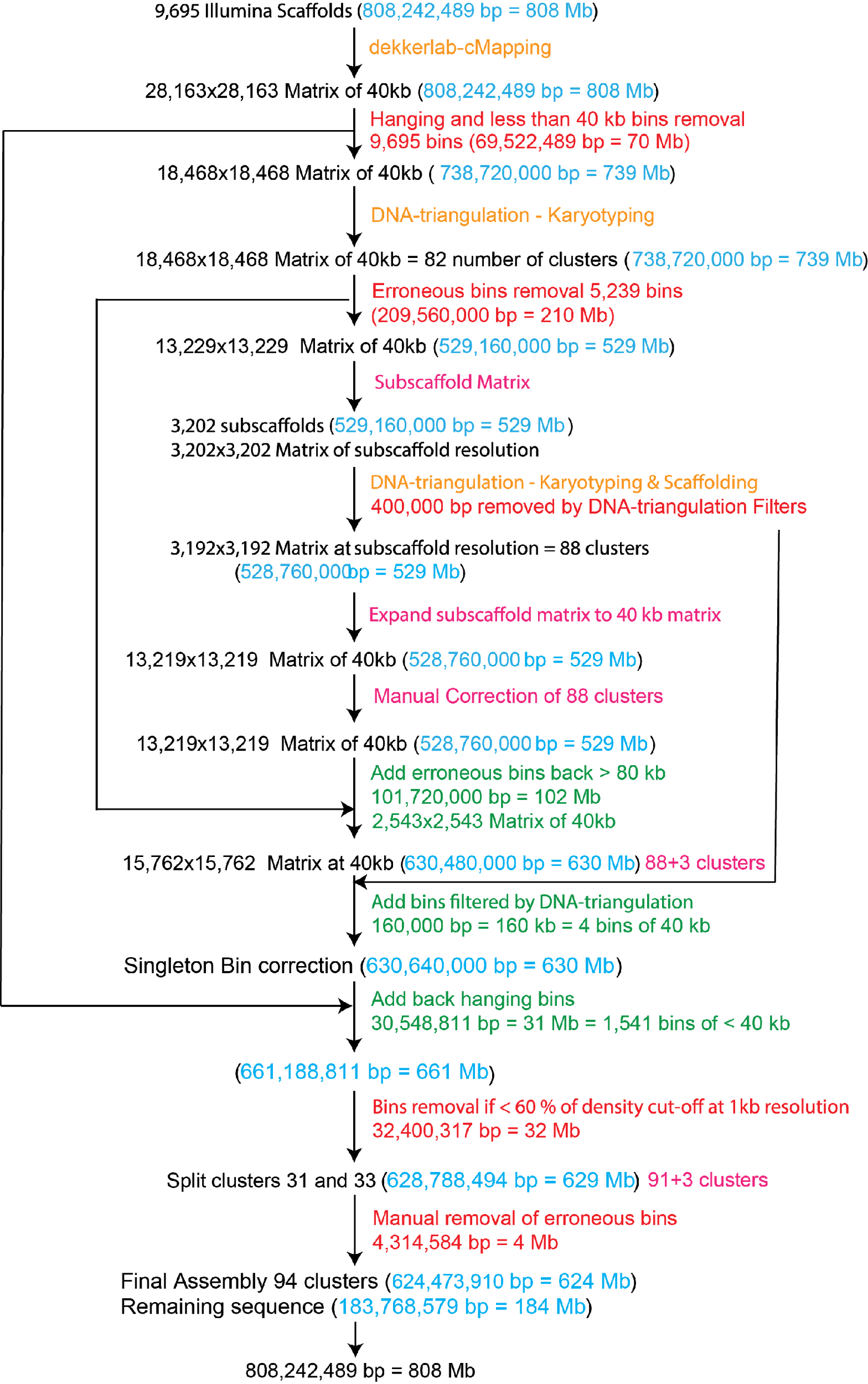

**Supplemental Figure S1**

**Assembly Flow Diagram**

Diagram of all steps used for Hi-C assisted chromosome scale assembly of the genome of *S. microadriaticum* (Smic1.0). The flow diagram indicates each step that is described in detail in the Methods section. Red text: steps where sequences were set aside; blue text: the total number of base pairs included in the assembly at the corresponding step; Orange: algorithmic steps using Hi-C read mapping Lajoie et al. Methods 2015 and the DNA triangulation method {Kaplan, 2013 #1182}; green text: number of base pairs placed back into the assembly; purple text: status of the Hi-C interaction matrix and number of chromosomes. A gapfilling step using PacBio reads was used to generate assembly version Smic1.1.

**SUPPLEMENTAL FIGURE S2**

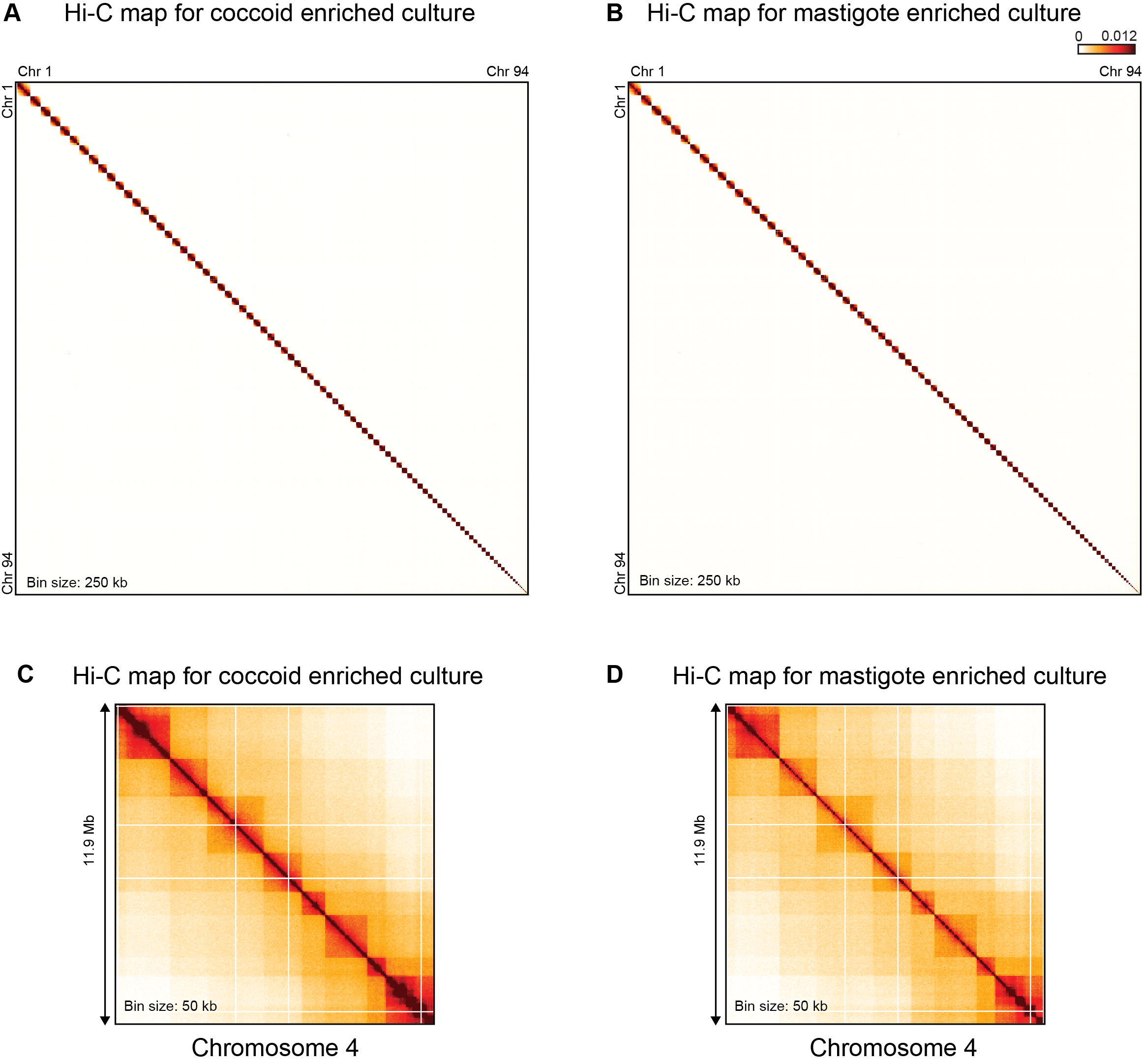

**Supplemental Figure S2**

Hi-C interaction maps obtained from coccoid-enriched and mastigote-enriched cultures reveal no obvious differences.

A. Genome-wide Hi-C interaction map for all 94 assembled chromosomes obtained with a coccoid-enriched culture. Bin size is 250 kb.
B. Genome-wide Hi-C interaction map for all 94 assembled chromosomes obtained with a mastigote-enriched culture. Bin size is 250 kb.
C. Hi-C interaction map for chromosome 4 obtained with a coccoid-enriched culture. Bin size is 50 kb.
D. Hi-C interaction map for chromosome 4 obtained with a coccoid-enriched culture. Bin size is 50 kb.

**SUPPLEMENTAL FIGURE S3**

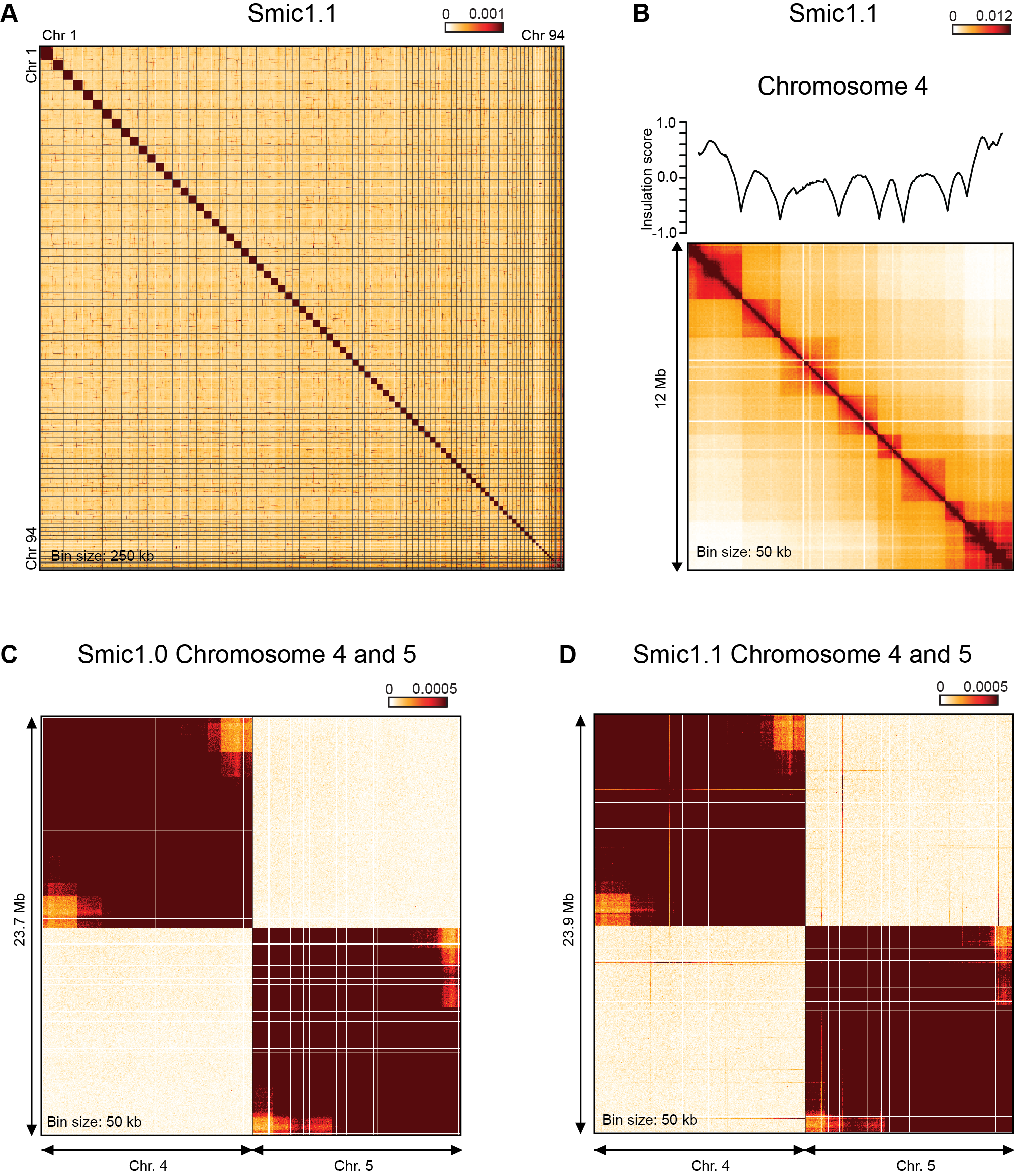

**Supplemental Figure S3**

Hi-C interaction maps for Smic1.1

A. Genome-wide Hi-C interaction map for all 94 assembled chromosomes. Bin size is 250 kb.
B. Hi-C interaction map for chromosome 4. Bin size is 50 kb. Plot on top of the heatmap represents the insulation profile (10 kb resolution, window size 500 kb; see Methods). Local minima in thus profiles indicate the locations of Hi-C domain boundaries.
C. Hi-C interaction map for Smic1.0 showing chromatin interactions for chromosomes 4 and 5 and interactions between chromosome 4 and 5. Binssize 50 kb. Color scale is chosen to highlight low-frequency inter-chromosomal interaction. Note the absence of prominent interactions between chromosome 4 and 5.
D. Hi-C interaction map for Smic1.1 showing chromatin interactions for chromosomes 4 and 5 and interactions between chromosome 4 and 5. Bin size 50 kb. Color scale is chosen to highlight low-frequency inter-chromosomal interaction. Note the presence of prominent lines of relatively frequent interactions between chromosome 4 and 5. These lines represent interactions between small regions on one chromosome with loci all along the other chromosomes. This indicates sequences may be present on multiple chromosomes.

**SUPPLEMENTAL FIGURE S4**

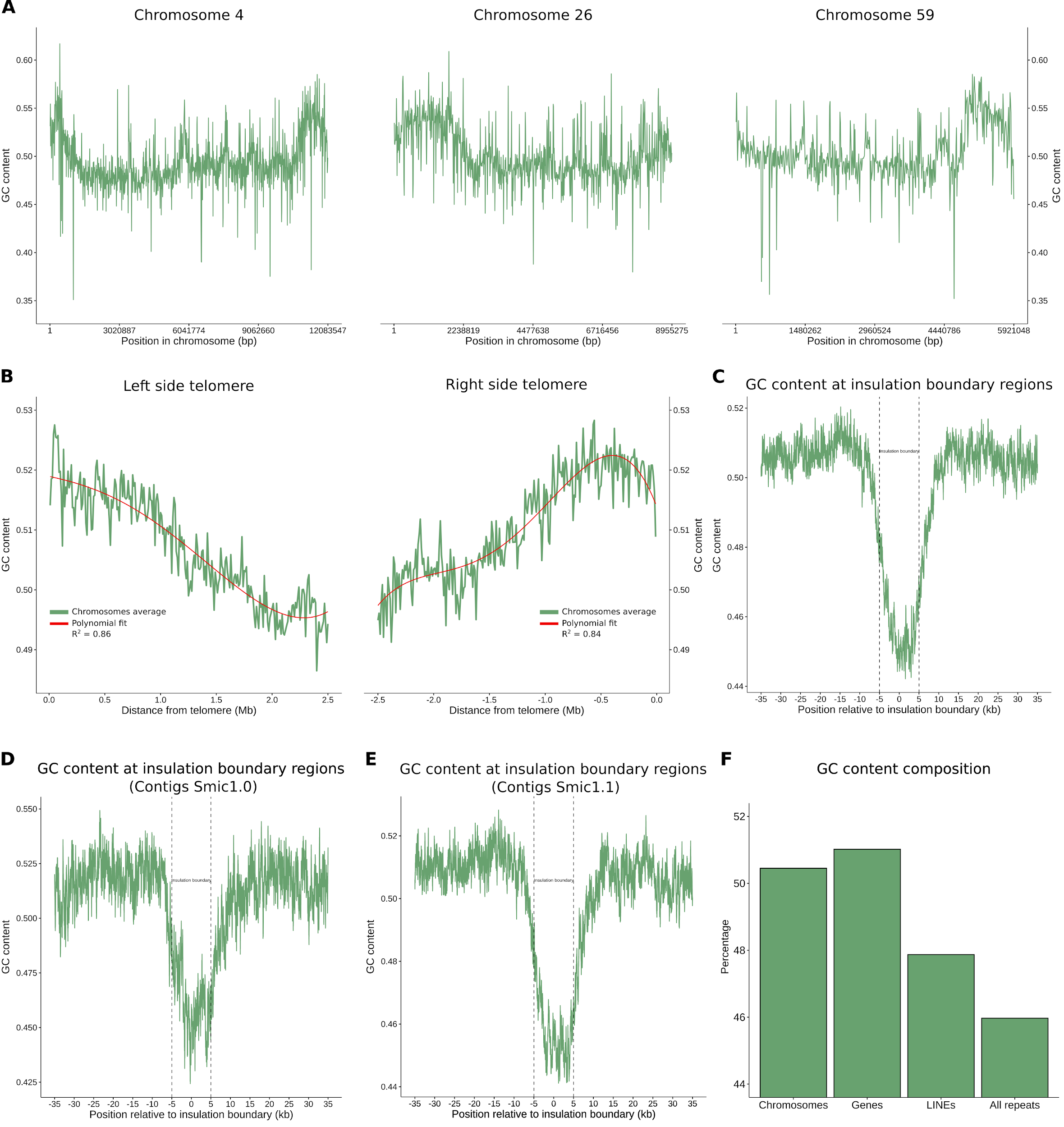

**Supplemental Figure S4**

**GC content along chromosomes and near telomeres for Smic1.1, and for Hi-C domain boundaries for Smic1.0 and Smic1.1**

A. GC content fluctuations along chromosomes 4, 26, and 59 measured in 10 kb windows for Smic1.1 assembly.
B. GC content along regions 2.5 Mb from telomeric ends, averaged for chromosomes of sizes of at least 5 Mb and measured in 10 kb windows for Smic1.1. GC content decreases as distance to telomeres increases.
C. GC content around insulation boundaries for Smic1.1. Values are averaged across all insulation boundaries in the genome for regions 30 kb upstream and downstream insulation boundaries and in 100 bp sliding windows. Dotted lines delimit insulation boundaries. A sharp decline in GC content is observed at insulation boundaries that define Hi-C domains.
D. GC content around insulation boundaries found in ungapped contigs for Smic1.0. This set of boundaries (each a 10 kb region) was located at least 10 kb from a contig end. Values are averaged across all insulation boundaries in the genome for 30 kb upstream and downstream insulation boundaries and in 100 bp sliding windows. Dotted lines delimit insulation boundaries. A sharp decline in GC content is observed at insulation boundaries that define Hi-C domains.
E. GC content around insulation boundaries found in ungapped contigs for Smic1.1. This set of boundaries (each a 10 kb region) was located at least 10 kb from a contig end. Values are averaged across all insulation boundaries in the genome for regions 30 kb upstream and downstream insulation boundaries and in 100 bp sliding windows. Dotted lines delimit insulation boundaries. A sharp decline in GC content is observed at insulation boundaries that define Hi-C domains.
F. GC content of chromosomes, genes, LINEs, and repetitive elements for Smic1.0. Suggesting that GC content is driven by genes.

**SUPPLEMENTAL FIGURE S5**

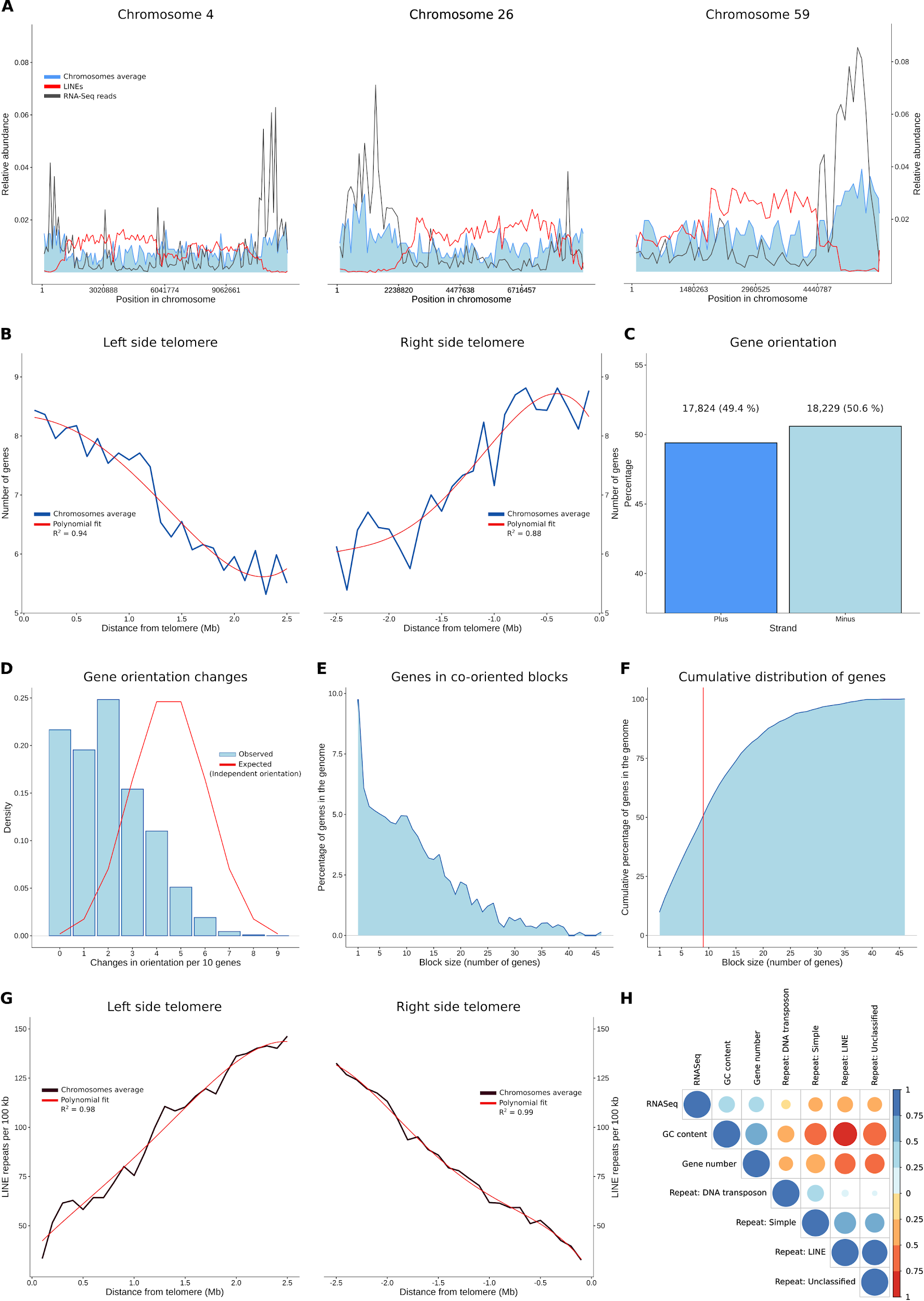

**Supplemental Figure S5**

**Gene and repetitive element distribution along chromosomes for Smic1.1**

A. Relative abundance of genes (blue), LINE repeats (red), and mapped RNASeq reads (grey) for chromosomes 4, 26, and 59.
B. Gene number along regions 2.5 Mb from telomeric ends, averaged for chromosomes of sizes of at least 5 Mb and measured in 100 kb windows. Gene number is observed to decrease as distance to telomeres increases.
C. Directionality of genes in the genome. A similar number of genes is found in both strands.
D. Frequencies of changes in gene orientation. Gene orientation changes defined as the occurrences of neighboring genes located in opposite strands and measured in sliding windows of 10 genes. Observed (blue) and assuming an equal and independent probability of gene orientation (red).
E. Distribution of genes in blocks of co-oriented genes.
F. Cumulative distribution of genes in blocks of co-oriented genes. 50 % of the genes are found in blocks of 9 or more co-oriented genes (red).
G. LINEs number along regions 2.5 Mb from telomeric ends, averaged for chromosomes of sizes of at least 5 Mb and measured in 100 kb windows. LINEs number is observed to increase as distance to telomeres increases.
H. Correlations between Gene number, GC content, RNASeq data, and Repeat types: LINE, DNA transposons, Simple, and Unclassified. Correlation coefficients are displayed as a color and size gradient between positive (blue) and negative (red) values. Correlations were done at 100 kb windows.

**SUPPLEMENTAL FIGURE S6**

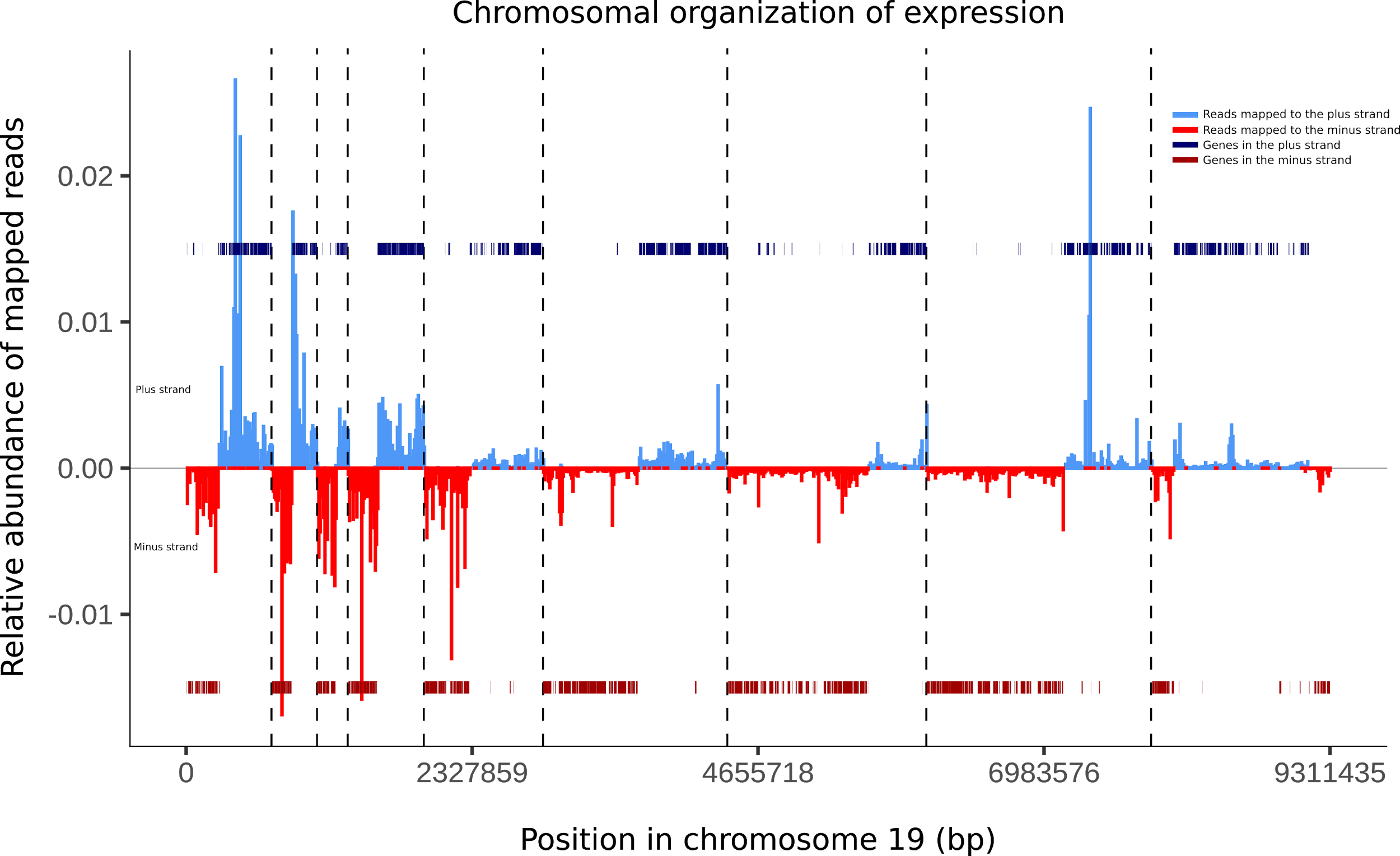

**Supplemental Figure S6**

**Chromosome 19 transcription and domain landscape for Smic1.1**

Indicated are: transcripts mapping to the plus strand (light blue), transcripts mapping to the minus strand (red), genes on the plus strand (dark blue), genes on the minus strand (dark red), and insulation boundaries as dotted vertical lines. A clear domainal gene block organization is observed and is delimited by insulation boundaries. Each domain is a pair of divergent gene blocks. Domain boundaries occur where gene blocks converge.

**SUPPLEMENTAL FIGURE S7**

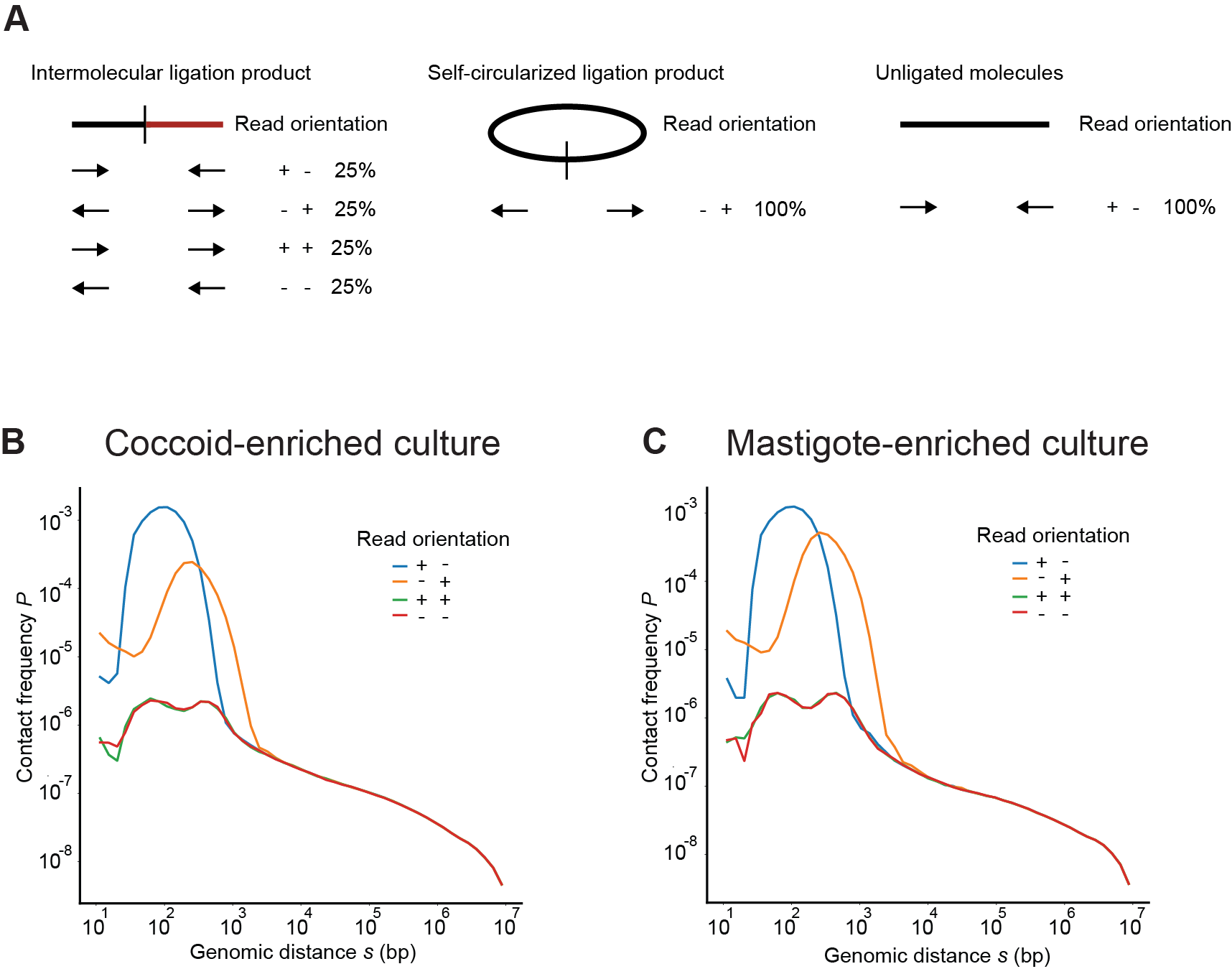

**Supplemental Figure S7**

***P*(*s*) plots for different Hi-C read orientations**

A. Different types of Hi-C ligation products. Ligation of two interacting restriction fragments can occur in 4 different fragment orientation (head-to-head, tail-to-tail, head-to-tail and tail-to-head). These 4 products can be identified by the Hi-C read orientation, as indicated, and are expected to occur with equal frequency (25% for each type). Self-circularized (partial) digestion products, or unligated linear fragments are also produced in Hi-C and are uninformative. These two products are characterized by the two Hi-C read orientations for each molecule: (-) and (+) for self-circularized molecules and (+) and (-) for unligated molecules.
B. Genome-wide *P*(s) plots for each of the 4 Hi-C read orientations for coccoid-enriched cultures. For distances below 2-3 kb uninformative Hi-C dominate, while for genomic distance over 2-3 kb all 4 read-orientation occur with equal frequency. Hi-C data was mapped against Smic1.0.
C. Genome-wide *P*(s) plots for each of the 4 Hi-C read orientations for mastigote-enriched cultures. For distances below 2-3 kb uninformative Hi-C dominate, while for genomic distance over 2-3 kb all 4 read-orientations occur with equal frequency. Hi-C data was mapped against Smic1.0.

**SUPPLEMENTAL FIGURE S8**

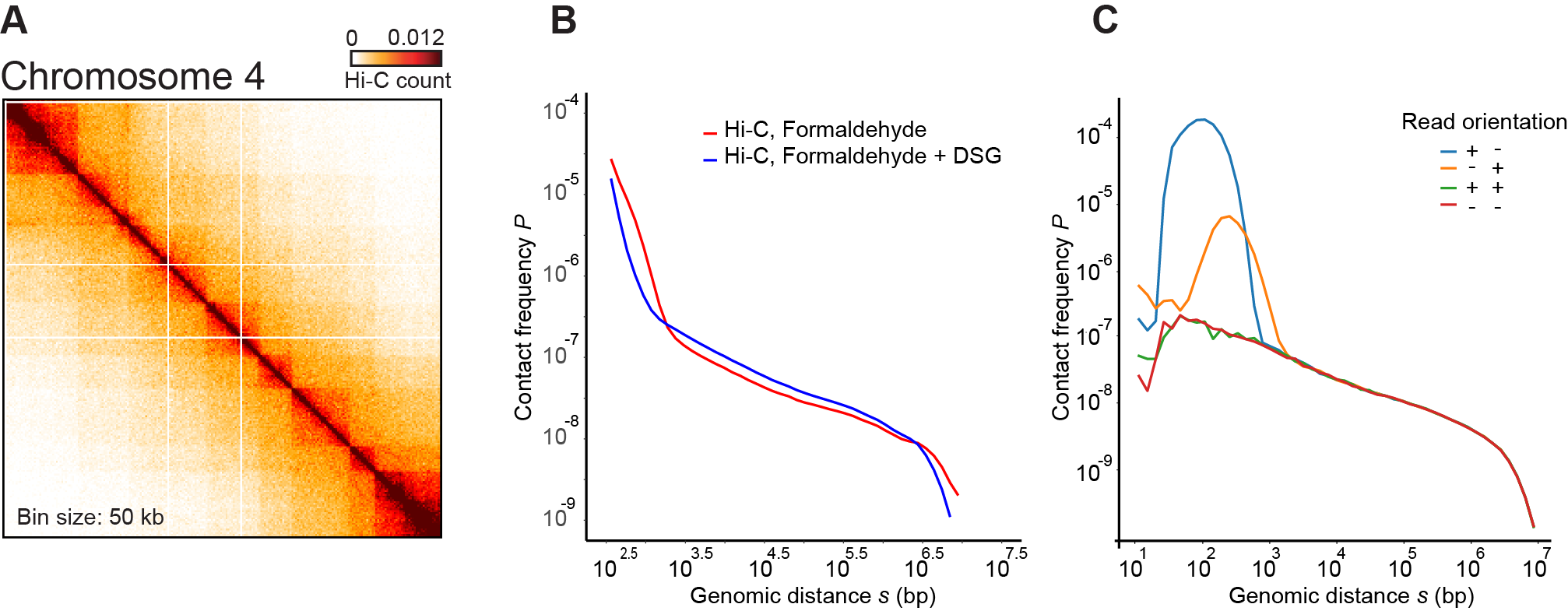

**Supplemental Figure S8**

**More extensive cross-linking does not greatly alter Hi-C maps and chromatin interaction frequencies**

A. Hi-C interaction map for chromosome 4 (bin size = 50 kb; Smic1.0) for coccoid enriched cultures fixed with formaldehyde and DSG.
B. Genome-wide contact frequency *P* versus genomic distance *s* for coccoid enriched cultures fixed with formaldehyde only, or with formaldehyde and DSG. *P*(*s*) displays three regimes (as in Figure 2A), and the middle section of the plots have similar small exponents. Hi-C data was mapped against Smic1.0.
C. Genome-wide contact frequency *P* versus genomic distance *s* for coccoid enriched cultures fixed with formaldehyde and DSG for each of the 4 possible orientation of ligation products. Ideally all 4 orientations display identical P(s). The + - and - + orientations deviate strongly for s < 2 kb and represent Hi-C artifacts (unligated ends and self-circularized molecules respectively. Hi-C data was mapped against Smic1.0. See Supplemental Figure S7 for additional details.

**SUPPLEMENTAL FIGURE S9**

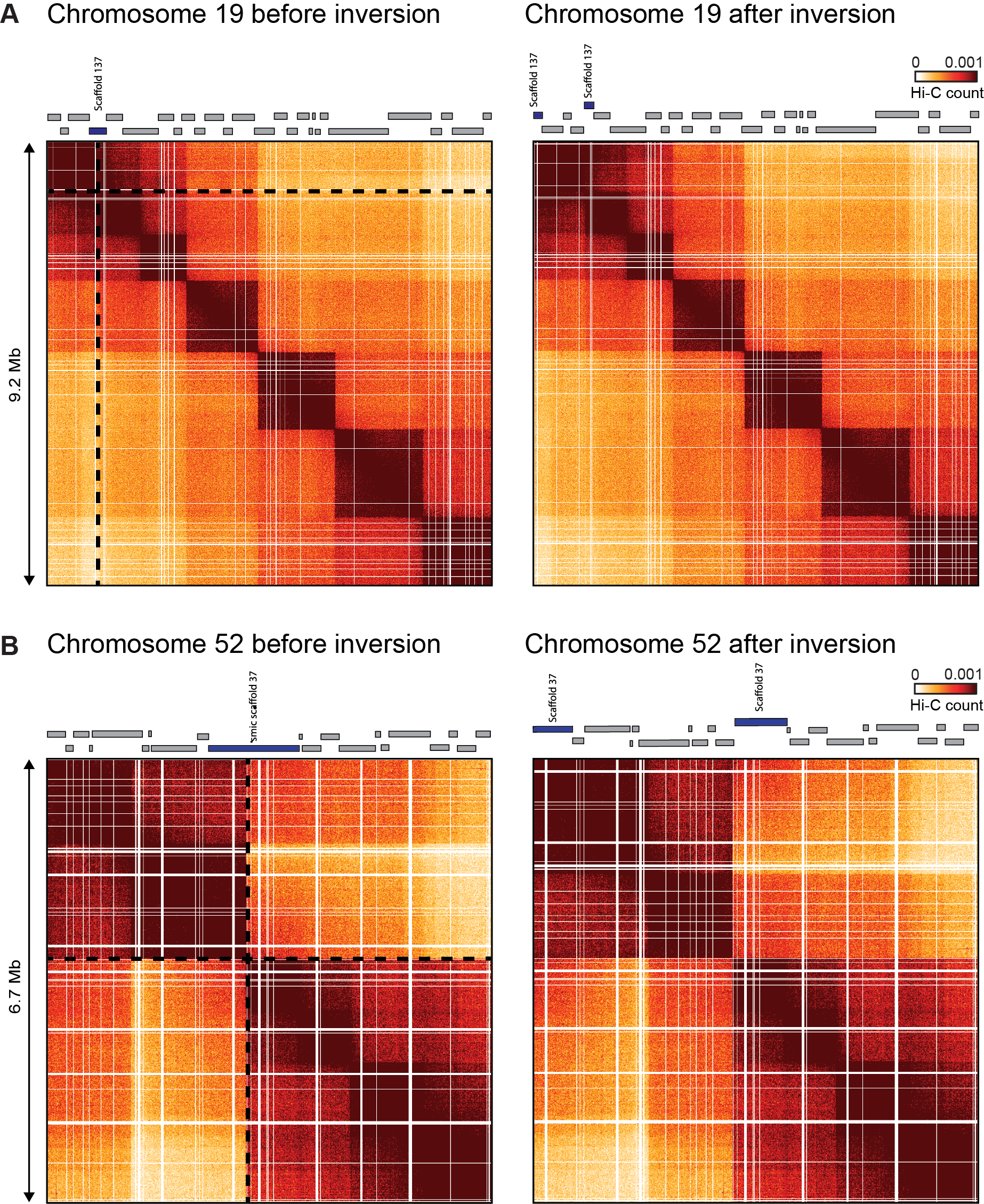

**Supplemental Figure S9**

**Examples of putative inversions in Smic1.0**

A. Left panel: Hi-C interaction map of chromosome 19 (bin size 10 kb). The rectangles on top of the heatmaps indicate positions of sub-scaffolds. The dotted lines indicate a potential misjoin, located within sub-scaffold 137 (blue rectangle). The Hi-C interaction pattern suggests that the section of position 1 up to the dotted line could be inverted. Right panel: as the left panel, but now with the sequence from position 1 up to the dotted line inverted. Inversion break sub-scaffold 137 in two pieces, as indicated by the two blue rectangles on top of the Hi-C interaction map.
B. Left panel: Hi-C interaction map of chromosome 52 (bin size 10 kb). The rectangles on top of the heatmaps indicate positions of sub-scaffolds. The dotted lines indicate a potential misjoin, located within sub-scaffold 37 (blue rectangle). The Hi-C interaction pattern suggest that the section of position 1 up to the dotted line could be inverted. Right panel: as the left panel, but now with the sequence from position 1 up to the dotted line inverted. Inversion breaks sub-scaffold 37 in two pieces, as indicated by the two blue rectangles on top of the Hi-C interaction map.

**SUPPLEMENTAL TABLES**

**SUPPLEMENTAL TABLE S1**

**See separate file for high resolution table**

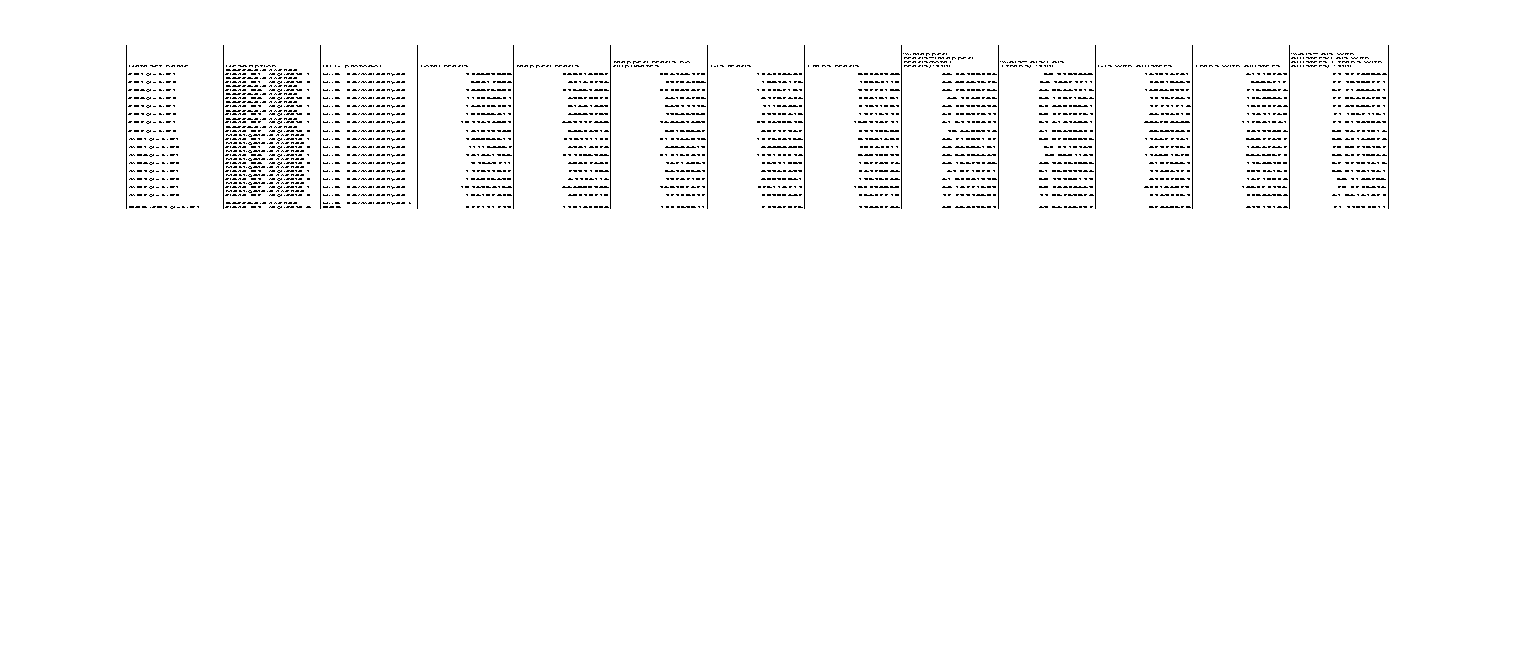

**Supplemental Table S1**

**Hi-C datasets for mastigote and coccoid-enriched cultures**

Hi-C data was generated for 4 clonal lines (D1, D3, D4 and D7). For each Hi-C libraries were prepared for mastigote and coccoid-enriched cultures. For each Hi-C dataset the following numbers are listed: total reads, mapped reads (to Smic1.0), number of cis interactions (interactions between loci located on the same assembled chromosome), trans interactions (interactions between loci located on different assembled chromosomes), percentage mapped reads, percent of all interactions that are in cis considering assembled chromosomes only, number of cis interactions including interactions within clusters 1-94, number of trans interactions including interactions that involve clusters 1-94, percent of all interactions that are in cis considering assembled chromosomes and clusters 1-94.

**SUPPLEMENTAL TABLE S2**

| **Assembly** | **Contig number** | **Scaffold N50** | **Contig N50** | **Gap %** | **Assembly size Scaffold/Contig** |
| --- | --- | --- | --- | --- | --- |
| Smic1.0 (Ns at all gaps) | 44997 | 8.44 Mb | 23.35 kb | 7.378 % | 624.696/580.637 Mb |
| Smic1.1 | 10628 | 8.47 Mb | 115.858 kb | 0.396 % | 626.052/624.259 Mb |

**Supplemental Table S2**

Summary statistics of Smic1.0 and Smic1.1, for the set of 94 chromosome-scale scaffolds.

**SUPPLEMENTAL TABLE S3**

**See separate file for high resolution table**

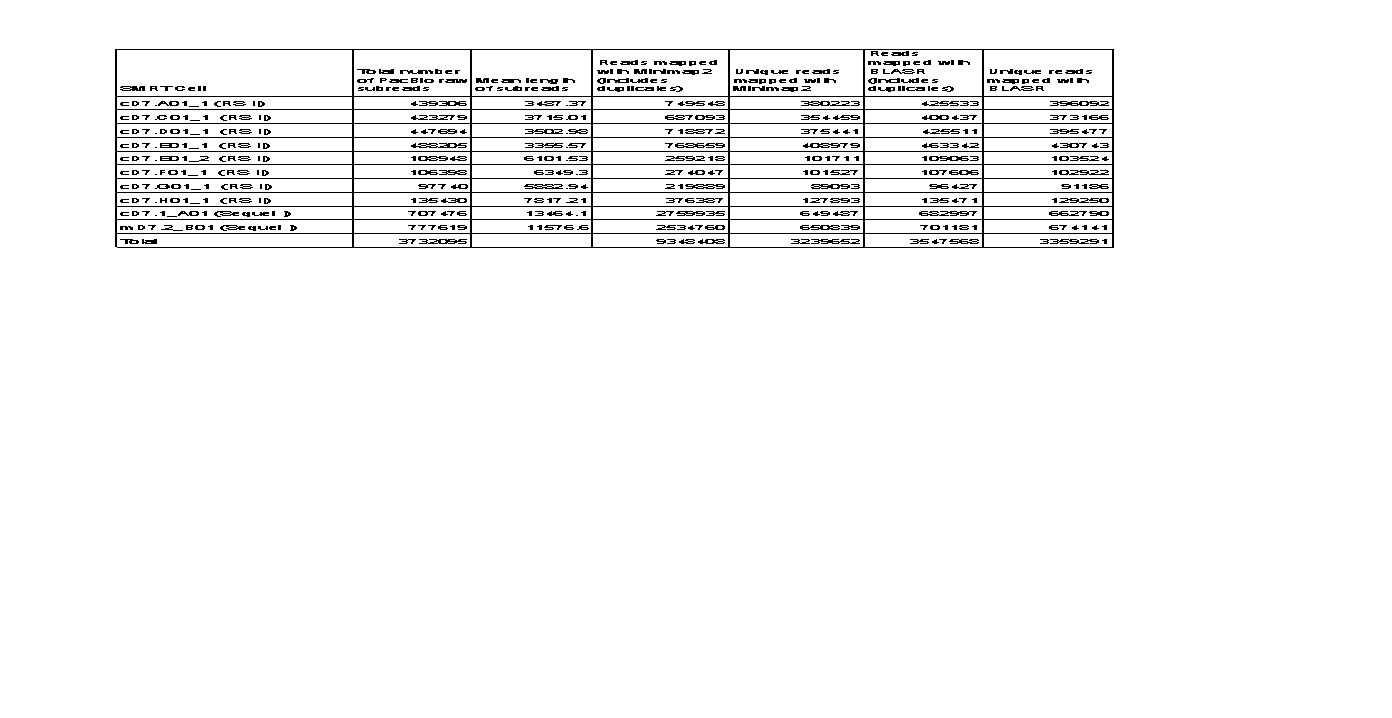

**Supplemental Table S3**

**PacBio datasets for mastigote and coccoid-enriched cultures for clone D7**

PacBio libraries were generated for mastigote-enriched cultures (mD7.2) and coccoid-enriched cultures (all other libraries). DNA was sequenced on a PacBio RSII instrument. For each library is listed: SMRT cell name, total number of raw subreads, mean length of the reads, number of reads mapped with Minimap2 including duplicates, number of reads mapped with Minimap2 excluding duplicates, number of reads mapped with BLASR including duplicates, and number of reads mapped with BLASR excluding duplicates.

**SUPPLEMENTAL DATA 1**

***Symbiodinium microadriaticum* genome v. 1.0 annotation file**

File containing gene coordinates, orientation, and annotations: BLAST hits against SwissProt and TrEMBL, InterPro classification, E.C. number, GO terms, and KEGG ontologies.

**SUPPLEMENTAL DATA 2**

***Symbiodinium microadriaticum* genome v. 1.0 gff3**

File containing gene tracks. File can be found here:

[https://drive.google.com/drive/folders/1IK7KtC2OI1UVEyYOjIDglAPkeh0JZEyC?usp=sharing](https://nam01.safelinks.protection.outlook.com/?url=https%3A%2F%2Fdrive.google.com%2Fdrive%2Ffolders%2F1IK7KtC2OI1UVEyYOjIDglAPkeh0JZEyC%3Fusp%3Dsharing&data=02%7C01%7CJob.Dekker%40umassmed.edu%7C64731683709c482849ea08d812d63c2f%7Cee9155fe2da34378a6c44405faf57b2e%7C0%7C0%7C637280058642542178&sdata=6MnHjl34NqpVzsG%2FQJlCeMU8ls2fyK%2Boc6aA%2BHm1mlU%3D&reserved=0" \o "Original URL: https://drive.google.com/drive/folders/1IK7KtC2OI1UVEyYOjIDglAPkeh0JZEyC?usp=sharing. Click or tap if you trust this link." \t "_blank)

**SUPPLEMENTAL DATA 3**

***Symbiodinium microadriaticum* genome v. 1.1 gff3**

File containing gene tracks. File can be found here:

[https://drive.google.com/drive/folders/1IK7KtC2OI1UVEyYOjIDglAPkeh0JZEyC?usp=sharing](https://nam01.safelinks.protection.outlook.com/?url=https%3A%2F%2Fdrive.google.com%2Fdrive%2Ffolders%2F1IK7KtC2OI1UVEyYOjIDglAPkeh0JZEyC%3Fusp%3Dsharing&data=02%7C01%7CJob.Dekker%40umassmed.edu%7C64731683709c482849ea08d812d63c2f%7Cee9155fe2da34378a6c44405faf57b2e%7C0%7C0%7C637280058642542178&sdata=6MnHjl34NqpVzsG%2FQJlCeMU8ls2fyK%2Boc6aA%2BHm1mlU%3D&reserved=0" \o "Original URL: https://drive.google.com/drive/folders/1IK7KtC2OI1UVEyYOjIDglAPkeh0JZEyC?usp=sharing. Click or tap if you trust this link." \t "_blank)

**SUPPLEMENTAL DATA 4-6**

**GO Enrichment analysis per chromosome for *Symbiodinium microadriaticum* genome v. 1.0**

GO enrichment analysis results from topGO. Files correspond to Biological Process, Molecular Function or Cellular Component categories.
