## Supplemental Table S1 for "Chromosome-scale assembly of the coral endosymbiont *Symbiodinium microadriaticum* genome provides insight into the unique biology of dinoflagellate chromosomes"

| Dataset name | Description | Hi-C protocol | Total reads | Mapped reads | Mapped reads no duplicates | Cis reads | Trans reads | %mapped reads=(mapped reads/total reads)*100 | %cis= cis/( cis +trans) *100 | Cis with clusters | Trans with clusters | %cis= cis with clusters/( cis with clusters + trans with clusters) *100 |
|---|---|---|---|---|---|---|---|---|---|---|---|---|
| cD1plus-R1 | Coccoid-enriched clone D1, replicate 1 | Hi-C, Formaldehyde | 423609808 | 238213087 | 205456470 | 125223532 | 80232938 | 56.23408205 | 60.9489358 | 154045731 | 51410739 | 74.97730833 |
| cD1plus-R2 | Coccoid-enriched clone D1, replicate 2 | Hi-C, Formaldehyde | 60317803 | 32152795 | 29705286 | 18843176 | 10862110 | 53.30564676 | 63.43374711 | 23018569 | 6686717 | 77.48980771 |
| cD3plus-R1 | Coccoid-enriched clone D3, replicate 1 | Hi-C, Formaldehyde | 433876802 | 246364306 | 222039372 | 122267184 | 99772188 | 56.78208765 | 55.06554216 | 150350997 | 71688375 | 67.71366521 |
| cD3plus-R2 | Coccoid-enriched clone D3, replicate 2 | Hi-C, Formaldehyde | 112063621 | 59870072 | 55105706 | 34787525 | 20318181 | 53.4250736 | 63.12871665 | 42467354 | 12638352 | 77.06525709 |
| cD4plus-R1 | Coccoid-enriched clone D4, replicate 1 | Hi-C, Formaldehyde | 153228204 | 81531339 | 65944446 | 41103362 | 24841084 | 53.20909393 | 62.33028631 | 47741713 | 18202733 | 72.39686721 |
| cD4plus-R2 | Coccoid-enriched clone D4, replicate 2 | Hi-C, Formaldehyde | 102836314 | 53689789 | 49636960 | 29920518 | 19716442 | 52.20897844 | 60.27870764 | 35295210 | 14341750 | 71.10671161 |
| cD7plus-R1 | Coccoid-enriched clone D7, replicate 1 | Hi-C, Formaldehyde | 1044545804 | 539447358 | 453534589 | 292590848 | 160943741 | 51.64420324 | 64.51345831 | 335703538 | 117831051 | 74.01939039 |
| cD7plus-R2 | Coccoid-enriched clone D7, replicate 2 | Hi-C, Formaldehyde | 131049930 | 63625913 | 60188637 | 30747957 | 29440680 | 48.5508943 | 51.08598322 | 36689332 | 23499305 | 60.95724015 |
| mD1plus-R1 | Mastigote-enriched clone D1, replicate 1 | Hi-C, Formaldehyde | 438083614 | 248441122 | 212455048 | 127623466 | 84831582 | 56.71089127 | 60.07080896 | 145577451 | 66877597 | 68.52153073 |
| mD1plus-R2 | Mastigote-enriched clone D1, replicate 2 | Hi-C, Formaldehyde | 111163367 | 59313273 | 53635519 | 33383308 | 20252211 | 53.35685181 | 62.2410459 | 37977962 | 15657557 | 70.80748487 |
| mD3plus-R1 | Mastigote-enriched clone D3, replicate 1 | Hi-C, Formaldehyde | 431351966 | 244286936 | 212162342 | 129122243 | 83040099 | 56.63285559 | 60.8601139 | 145601670 | 66560672 | 68.62748055 |
| mD3plus-R2 | Mastigote-enriched clone D3, replicate 2 | Hi-C, Formaldehyde | 94659711 | 50327532 | 45715064 | 26941089 | 18773975 | 53.16679236 | 58.93262886 | 31076664 | 14638400 | 67.97904516 |
| mD4plus-R1 | Mastigote-enriched clone D4, replicate 1 | Hi-C, Formaldehyde | 147844827 | 79941283 | 65430634 | 39952599 | 25478035 | 54.0710721 | 61.06099935 | 44505472 | 20925162 | 68.01931951 |
| mD4plus-R2 | Mastigote-enriched clone D4, replicate 2 | Hi-C, Formaldehyde | 105026590 | 54425115 | 49737107 | 30090851 | 19646256 | 51.82031998 | 60.49980149 | 34027084 | 15710023 | 68.4138786 |
| mD7plus-R1 | Mastigote-enriched clone D7, replicate 1 | Hi-C, Formaldehyde | 1045963163 | 555800936 | 458407574 | 276113714 | 182293860 | 53.13771609 | 60.23323559 | 322135079 | 136272495 | 70.2726345 |
| mD7plus-R2 | Mastigote-enriched clone D7, replicate 2 | Hi-C, Formaldehyde | 105107328 | 50240710 | 47428247 | 20900537 | 26527710 | 47.79943602 | 44.06769873 | 24592264 | 22835983 | 51.85151372 |
| DSG-cD4plus-R1 | Coccoid-enriched clone D4, replicate 3 | Hi-C, Formaldehyde+ DSG | 277141749 | 140132003 | 122269811 | 72937076 | 49332735 | 50.56329604 | 59.65256297 | 87350678 | 34919133 | 71.44092011 |
