## Supplemental S3 for "Chromosome-scale assembly of the coral endosymbiont *Symbiodinium microadriaticum* genome provides insight into the unique biology of dinoflagellate chromosomes"

| SMRT Cell | Total number of PacBio raw subreads | Mean length of subreads | Reads mapped with Minimap2 (includes duplicates) | Unique reads mapped with Minimap2 | Reads mapped with BLASR (includes duplicates) | Unique reads mapped with BLASR |
|---|---|---|---|---|---|---|
| cD7.A01_1 (RS II) | 439306 | 3487.37 | 749548 | 380223 | 425533 | 396092 |
| cD7.C01_1 (RS II) | 423279 | 3715.01 | 687093 | 354459 | 400437 | 373166 |
| cD7.D01_1 (RS II) | 447694 | 3502.98 | 718872 | 375441 | 425511 | 395477 |
| cD7.E01_1 (RS II) | 488205 | 3355.57 | 768659 | 408979 | 463342 | 430743 |
| cD7.E01_2 (RS II) | 108948 | 6101.53 | 259218 | 101711 | 109063 | 103524 |
| cD7.F01_1 (RS II) | 106398 | 6349.3 | 274047 | 101527 | 107606 | 102922 |
| cD7.G01_1 (RS II) | 97740 | 5882.94 | 219889 | 89093 | 96427 | 91186 |
| cD7.H01_1 (RS II) | 135430 | 7817.21 | 376387 | 127893 | 135471 | 129250 |
| cD7.1_A01 (Sequel I) | 707476 | 13464.1 | 2759935 | 649487 | 682997 | 662790 |
| mD7.2_B01 (Sequel I) | 777619 | 11576.6 | 2534760 | 650839 | 701181 | 674141 |
| Total | 3732095 | | 9348408 | 3239652 | 3547568 | 3359291 |
